# Brain dynamics of mental manipulation

**DOI:** 10.1101/2025.10.20.683458

**Authors:** David R. Quiroga-Martinez, Tianyu He, Gemma Fernández-Rubio, Leonardo Bonetti, Alejandro O. Blenkmann, Tor Endestad, Anne-Kristin Solbakk, Torstein R. Meling, Martin Fabricius, Olivia Kim-McManus, Jon T. Willie, Peter Brunner, Mohammad Dastjerdi, Jack J. Lin, Robert T. Knight

## Abstract

Humans effortlessly juggle their internal thoughts, but the neuronal dynamics that support mental manipulation are largely unknown. Leveraging the high spatiotemporal fidelity of intracranial recordings in humans (N = 30), we provide evidence that mental sound manipulation involves the inhibition of sensory cortex and the coordinated engagement of memory and control networks. This modulation manifests in two ways. First, there is a shift in the balance between faster (> 30 Hz) and slower (< 30 Hz) dynamics in primary and secondary auditory areas, suggesting a decrease in local excitability. Second, there is a distributed increase in oscillatory synchrony (6-10 Hz), which predicts imagery vividness and task performance. This evidence points to a key role of local excitability and inter-areal synchrony in the manipulation of thought.

## 1. Introduction

Imagine creating a song for someone you cherish. You may not be a musician, but you take inspiration from your favorite songs. In your mind, you begin stitching lyrics together, crafting a simple melody, and rearranging bits and pieces until you end up with a catchy tune. The **mental manipulation** of our inner thoughts, whether musical or of any kind, is essential for capacities such as reasoning, planning, problem-solving, and creativity, which are the hallmark of human cognition. Yet, the neuronal mechanisms underlying the manipulation of thought remain largely unknown.

The temporary storage and manipulation of information is known as **working memory** (WM) (1). This ability involves widespread brain networks, with control linked to frontoparietal areas (2), content representation linked to sensory areas (3, 4), and long-term memory contributions linked to medial temporal structures (5–7). Functional and structural connections among these regions are crucial for WM (8, 9).

WM entails two operations: **maintenance** (keep as is) and **manipulation** (transform into something else). However, the latter has remained surprisingly understudied. Typically, WM experiments manipulate task demands by including distractors or increasing the number of items to remember (e.g., 10, 11), with only a few studies directly investigating the transformation of mental objects (12–18). How does the brain implement mental manipulation?

Information in WM is thought to be maintained in persistent activity (19) or patterns of synaptic strength (20). In both cases, the neural code must be reconfigured to represent the transformed information. Manipulation elicits larger hemodynamic responses in frontoparietal cortex (12, 14), indicating recruitment of neural resources for cognitive control. Manipulation improves with parietal theta stimulation (13) and enhances frontal midline theta (1 - 8 Hz) dynamics in scalp electroencephalography (EEG) (16, 17). This suggests that neuronal dynamics play an essential role in manipulation, but the exact mechanisms by which they do so remain unclear.

We leveraged the high spatiotemporal fidelity of intracranial EEG (iEEG) to investigate the brain dynamics of mental manipulation. Using an auditory WM task (13, 21), we first asked how manipulation changes patterns of neural activity across frequency bands and brain areas as an index of local excitability. We then tested whether manipulation modulates network synchrony as an indicator of inter-areal communication, and evaluated the potential role of phase-amplitude coupling between slow and fast neuronal dynamics. We provide evidence that manipulation involves the inhibition of primary and secondary sensory cortex and the engagement of memory and control networks via neural synchrony.

## 2. Results

A group of presurgical epilepsy patients (N = 30, 16 female, age = 37.32 +/− 11.91 SD) performed an auditory WM task. On each trial, participants listened to a three-note melody and were asked to vividly imagine it immediately after during a silent delay (2 s) (**Fig. 1A**). We encouraged vivid imagery to constrain retention strategies (22) and maximize active mental states. There were two conditions. In the **recall** block, participants imagined the melodies as presented, while in the **manipulation** block, they imagined them in reverse. Subsequently, participants answered whether a test melody matched the original sample (recall) or its inverted version (manipulation). Two melodies were to be imagined in the task (mel1: A-C#-E, mel2: E-C#-A).

**Fig 1.**
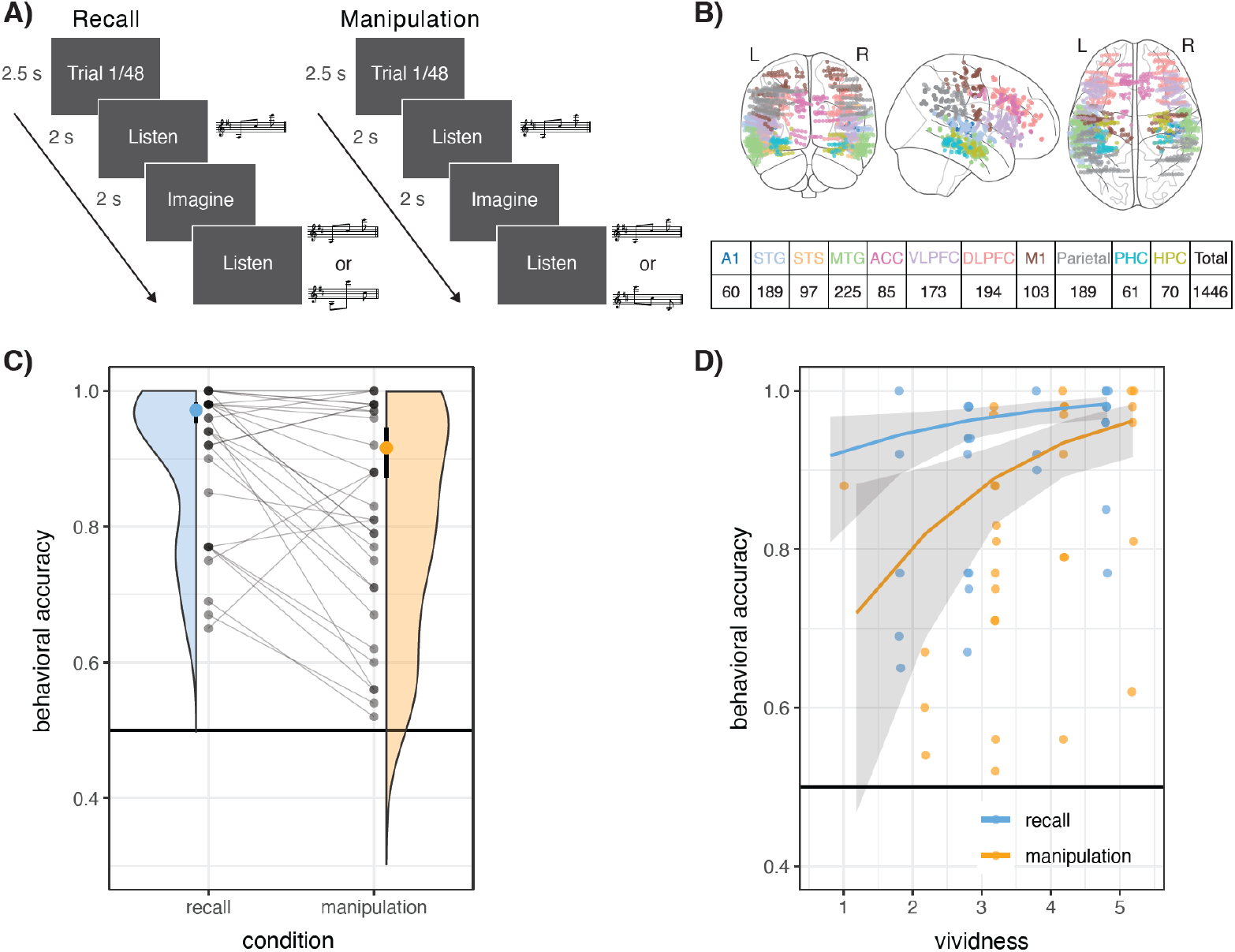
Experimental design and behavior. **A)** Participants completed an auditory working memory task where they heard a short melody, were asked to vividly imagine it, and then compare it with a test melody. In the recall block, participants imagined the melody as presented, while in the manipulation block they imagined it backwards (e.g., A-C#-E becomes E-C#-A). **B)** We recorded intracranial EEG from 11 regions of interest (see **Fig. S1** for electrodes per region). **C)** Participants performed above chance and were better during recall than manipulation. **D)** Task performance was predicted by ratings of imagery vividness in both blocks. Error bars and shaded areas represent 95% confidence intervals, as obtained from mixed-effects logistic regression.

Task performance was high and was lower (χ^2^(1) = 68.39, *p* < .001, OR = 0.318, CI = [0.204– 0.496]) for manipulation (91.6%, CI = [87.19% - 94.58%]) than recall (97.17%, CI = [95.34% −98.29%]) (**Fig. 1C & table S1**). After each block, participants rated imagery vividness from 1 (not vivid at all) to 5 (extremely vivid). These ratings were not significantly different between conditions (χ^2^(1) = 0.426, *p* = .514, OR = 0.68, CI = [0.21 - 2.14]) (**table S2**), with most participants rating three or above (recall: 25/30 = 83.33%, manipulation: 26/30 = 86.66%). Moreover, vividness predicted task performance (χ^2^(1) = 7.72, *p* = .005, OR = 1.61, CI = [1.16 - 2.25]), suggesting participants used imagery for WM control (**Fig. 1D**). We found no significant association of music training with task performance (χ^2^(1) = 3.08, *p* = .079, OR = 2.29, CI = [0.95 – 5.49]) or vividness ratings (χ^2^(1) = 1.45, *p* = .233, OR = 5.11, CI = [0.31 – 82.6]). We selected correct trials for further analyses.

### 2.1 Distinct spectrotemporal profiles during WM

We chose 11 brain regions of interest (ROIs) based on the WM literature (**Fig. 1B & Fig. S1**): Primary auditory cortex (A1), superior temporal gyrus (STG), superior temporal sulcus (STS), middle temporal gyrus (MTG), anterior cingulate (ACC), ventrolateral prefrontal cortex (VLPFC), dorsolateral prefrontal cortex (DLPFC), motor cortex (M1), hippocampus (HPC), parahippocampal (PHC), and parietal cortex. For statistical analyses, we fitted Bayesian mixed-effects models to z-scored (log) power values with participant-wise random effects (**table S3**). We did this separately for each region, time point, and seven canonical frequency bands: Low-theta (1-4 Hz), high-theta (4-8 Hz), alpha (8-12 Hz), low-beta (12-20 Hz), high-beta (20-30 Hz), low-gamma (30-70 Hz), and high-gamma (70-160 Hz).

We first characterized the mean spectrotemporal dynamics of each region by examining models with an intercept term as the only predictor. Compared to baseline, A1 and STG generated strong responses to each perceived sound in the gamma and high-gamma range (**Fig. 2B-C, Fig S2-S4**). In lower frequencies (low-theta, high-theta, alpha, low-beta, high-beta), there was an initial bump of activity at melody onset, quickly followed by suppression below baseline. All frequency bands tended towards baseline levels during the delay period. This pattern of response, referred to as **sensory profile**, is mainly driven by the presence or absence of sounds.

**Fig 2.**
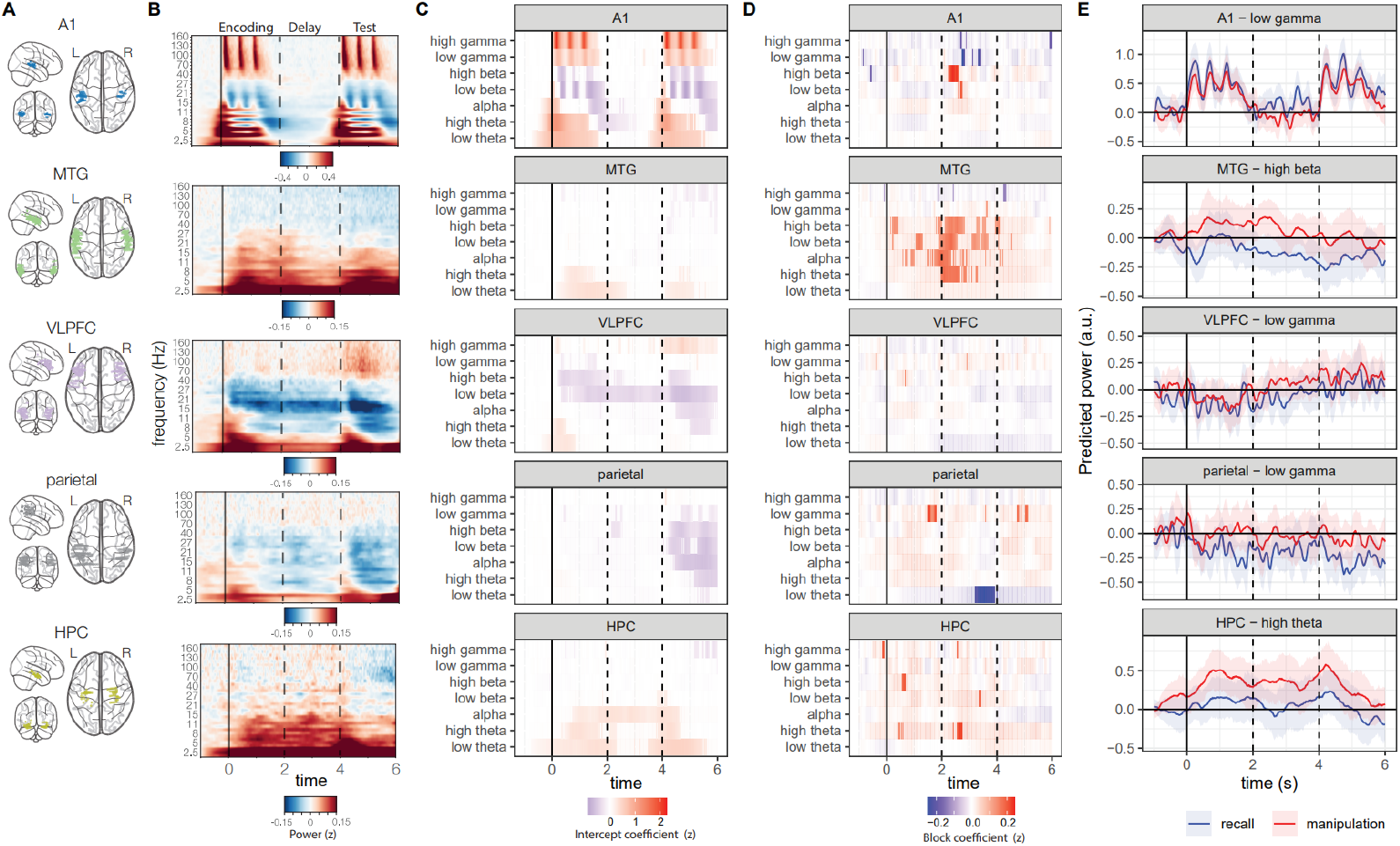
Time-frequency analyses. **A)** Across brain areas, **B)** average time-frequency representations revealed different spectral profiles in neuronal dynamics (see **Fig. S2** for all regions). Regression coefficients show these dynamics were modulated by **C)** the task compared to baseline and by **D)** manipulation at different frequency bands (**Fig. S3**). **E)** Model predictions for representative frequency bands further depict differences between conditions (**Fig. S4**). Highlighted segments mark time points with conclusive evidence. Shaded areas indicate 95% highest density intervals. Solid and dashed vertical lines mark the onset of encoding, delay and test periods.

In frontoparietal regions (VLPFC, DLPFC, ACC, M1, parietal), we observed a similar task-related suppression of lower frequencies (alpha, low-beta, high-beta), with a transient low- and high-theta increase at trial onset (**Fig. 2B-C & Fig S2**). After the start of the test melody, high-gamma increased and high- and low-beta further decreased, likely reflecting decision making and motor responses. Furthermore, across the whole trial, low-theta activity remained above baseline, while VLPFC and DLPFC showed persistent high-gamma activity, possibly reflecting the maintenance of information in WM. We refer to this response pattern as **dorsal profile**.

Finally, STS, MTG, HPC, and PHC exhibited a task-related increase in lower frequencies (low-theta, high-theta, alpha) with a tendency to decrease in higher frequencies (**Fig. 2B-C & Fig S2**). This response pattern, which we refer to as **ventral profile**, was sustained during the trial, ending at the beginning of the test melody.

A common theme is that the enhancement of higher frequencies is accompanied by the suppression of lower frequencies and vice versa, likely reflecting changes in excitation / inhibition balance (23). Importantly, dorsal and ventral areas appear to exhibit opposite dynamics, with higher frequencies dominating dorsal areas and lower frequencies dominating ventral areas.

### 2.2. Manipulation modulates brain dynamics across the frequency spectrum

Next, we investigated how mental manipulation changes neuronal dynamics. We included “block” (manipulation vs. recall) and its random slope as predictors in statistical models, which we compared using Deviance Information Criteria (DIC) and model probabilities (or weights) derived from them (**table S3**). We consider evidence for an effect as conclusive if the 95% highest-density interval (HDI) around the block coefficient excludes 0 and the cumulative probability of models including block as a predictor is at least 0.95.

Manipulation shifted the balance between slower and faster dynamics in primary and secondary auditory areas. In A1, it enhanced low-beta and high-beta power while decreasing low-gamma and high-gamma power during the delay period (**Fig. 2D**). In the MTG, there was a robust increase in theta, alpha, low-beta, and high-beta power across the trial, with sparse decreases in high-gamma.

Other regions exhibited more sparse modulations of activity. For example, short-lived enhancements were seen in HPC across the spectrum, most notably in high-theta (**Fig. 2D**). In the parietal cortex, low-gamma was enhanced, and low-theta was briefly suppressed by manipulation. Low-gamma increases were also seen in VLPFC, and low-beta enhancements were detected in PHC (See **Fig. S3** for full report). Taken together, these results suggest mental manipulation primarily enhances inhibition in primary and secondary auditory areas, as reflected in the balance between faster and slower neuronal dynamics.

### 2.3. Patterns of neuronal dynamics distinguish manipulation from recall

Apart from changes in the mean activity of brain regions, manipulation may result in different activity patterns across neuronal populations. To investigate this possibility, we used multivariate analyses to assess whether context information (recall vs. manipulation) was encoded in patterns of neuronal dynamics. We used logistic regression to decode block identity from power estimates in different frequency bands, for each subject and the encoding and delay periods separately. We did this both within each ROI (local decoding) and pooling together all the ROIs available for the subject (global decoding). In these models, signals at each channel and time-point were entered as features or predictors. For each period and region separately, we used Bayesian mixed-effects models to determine whether decoding accuracies were conclusively different from chance (0.5). We entered frequency band and subject as grouping variables for random intercepts (**table S4**). To characterize decoding accuracies across bands, we estimated intercept-only models and inspected random-effect coefficients for interpretation.

During the encoding period, global decoding was above chance for most frequency bands, with peak accuracies in high-gamma and low-theta (**Fig. 3A**). In turn, local decoding was above-chance for STG high-gamma and high-theta, MTG high-theta, and A1 low-theta (**Fig. 3B**). During the delay period, above-chance global decoding was detected for high-gamma, high-theta, low-theta, and alpha signals, whereas local decoding was above-chance in MTG high-gamma and M1 low-theta. Consistent with univariate analyses, these results suggest manipulation mainly changes patterns of activity in primary and secondary auditory areas, with theta and high-gamma dynamics showing the clearest effects.

**Fig 3.**
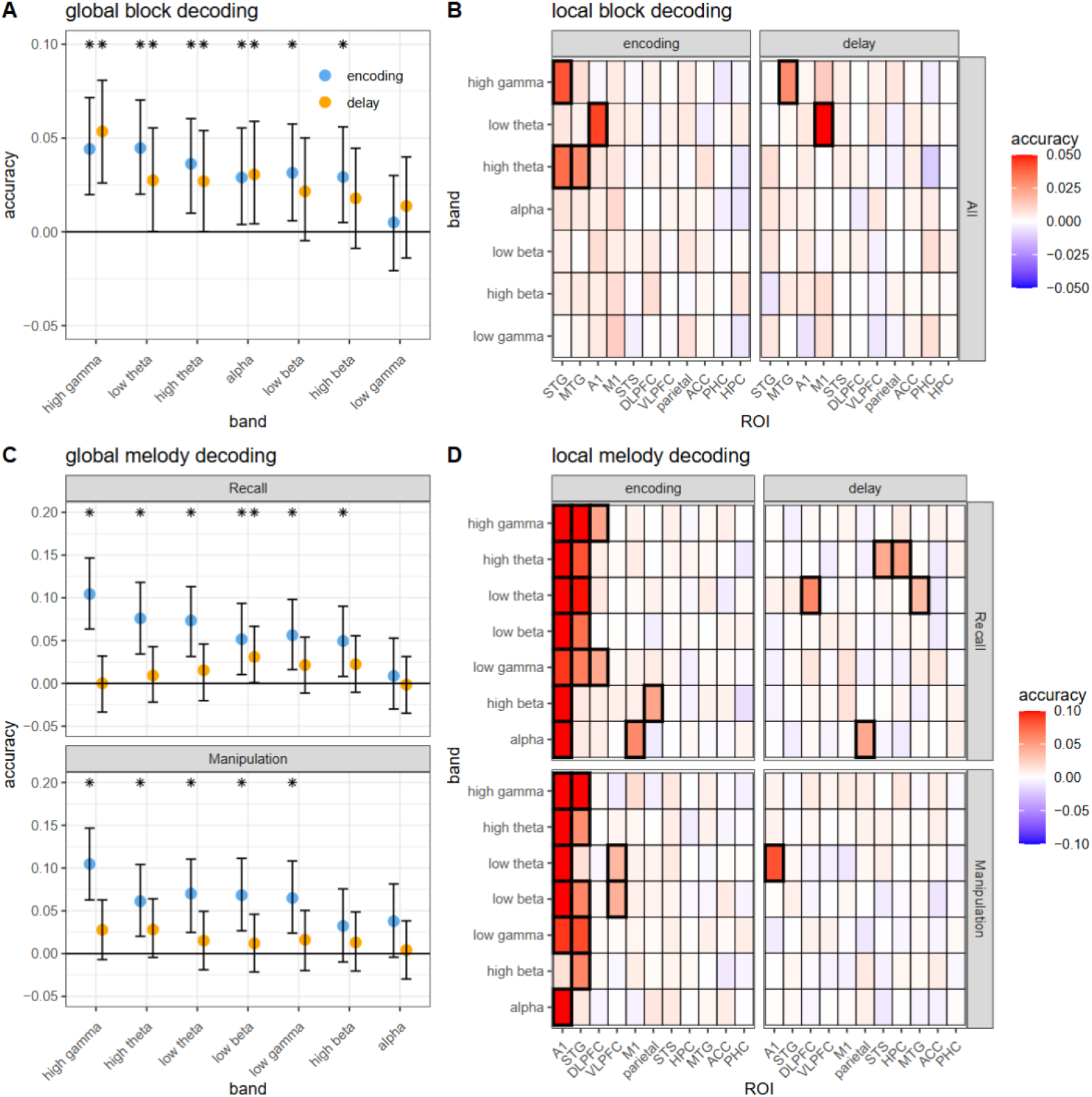
Decoding. Information about context (manipulation vs. recall) was successfully decoded from encoding and delay **A)** global and **B)** local dynamics. In contrast, **C)** global decoding of melody identity was conclusively above chance in the encoding but not the delay period. **D)** Local decoding indicates auditory and prefrontal areas were the main carriers of melody information during encoding. Accuracies are presented relative to chance (0.5). Error bars mark 95% HDI. Asterisks and highlighted squares indicate regions and frequency bands whose 95% HDI excluded chance level.

### 2.4. Neuronal dynamics carry information about auditory objects

Next, we investigated whether neuronal dynamics also carry information about the perceived and imagined melodies. We used the same decoding approach to predict melody identity (mel1 vs. mel2) from neuronal dynamics for each block separately. During the encoding period, global decoding was successful for most frequency bands, with a peak in high-gamma (**Fig. 3C**). In turn, local decoding accuracy was highest in A1 and STG, but was also above-chance in DLPFC, M1, and parietal for the recall block, and VLPFC and M1 for manipulation (**Fig. 3D**).

In contrast, delay-period information was more difficult to retrieve. We found above-chance accuracy for low beta global decoding only. Nevertheless, local decoding suggested above-chance performance for DLPFC and MTG low-theta, STS and HPC high-theta, and parietal alpha during recall, together with A1 low-theta during manipulation. However, due to the lack of clear global decoding effects, the contribution of each of these regions needs to be taken with caution.

In sum, while the identity of perceived sounds is readily available in auditory and prefrontal areas, especially in the high-gamma range, information about imagined sounds is more difficult to retrieve from conventional iEEG metrics, especially during manipulation. This points to a fine-grained encoding of imagined auditory objects in neuronal populations.

### 2.5. Manipulation enhances directed high-theta / low-alpha synchrony

Lower-frequency dynamics are proposed to facilitate communication between brain regions through synchrony (24, 25). Given that manipulation requires the exchange of representation and control signals, we hypothesized it would lead to enhanced low-frequency inter-areal synchrony. We estimated coherence-based directed functional connectivity between pairs of electrodes in different ROIs during the silent delay for each condition separately. We subtracted the pretrial baseline to obtain delay-specific connectivity. We obtained the (log absolute) imaginary part of coherence (imCOH) (26) for frequencies between 2 and 30Hz, in 2 Hz bins. For each bin, we fitted and compared several Bayesian mixed-effects models in which both subject and connection (i.e., pair of ROIs) were included as grouping variables for random intercepts and slopes (**table S5**). We used the same criteria described above to identify conclusive evidence.

Across ROIs, manipulation enhanced synchrony in the theta and alpha range (6-10Hz), with a peak in the 8-10Hz frequency bin (β = 0.063, 95% HDI = [0.02 - 0.10]) (**Fig 4A-B**). Random-effect coefficients at the peak revealed conclusive evidence for modulations between auditory (A1, STG, STS, MTG) and association (DLPFC, HPC, VLPFC, ACC, M1) areas (**Fig. 4D-E**) and conclusive effects for STG-A1, VLPFC-HPC, and parietal-M1.

**Fig 4.**
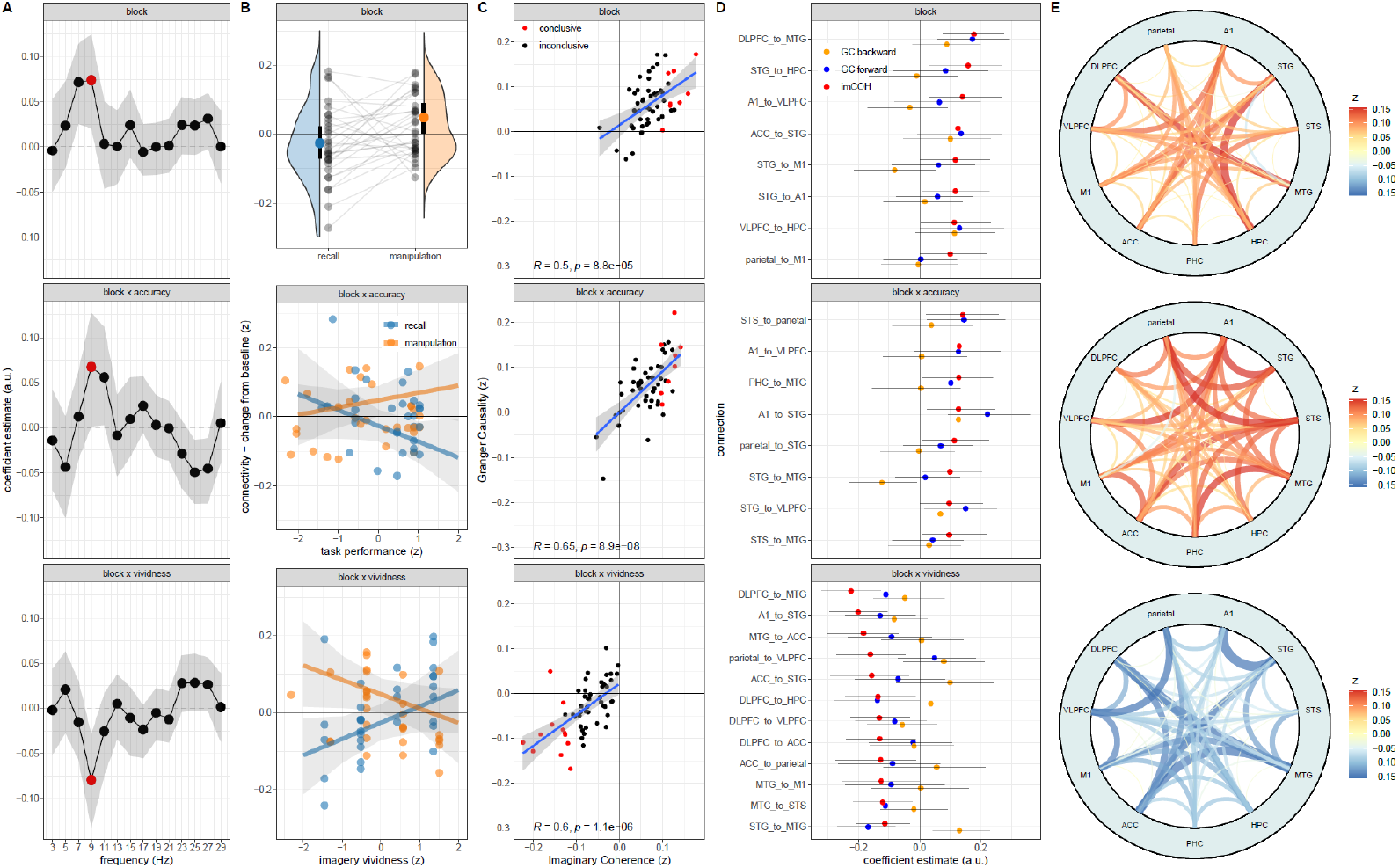
Connectivity. **A)** Regression coefficients reveal a brain-wide enhancement of high-theta and low-alpha (6-10 Hz) imaginary coherence (imCOH) during manipulation (top; block effect) and low-alpha interactions with (z-scored) task performance (i.e., accuracy; middle) and imagery vividness (bottom). Red dots highlight frequencies with conclusive effects. **B)** The effect of block and its interaction with task performance and vividness are also shown in model predictions for the 8-10 Hz frequency bin (i.e., posterior medians) overlaid over connectivity values averaged across all electrode pairs separately for each participant. For task performance, data is shown after removing effects of imagery vividness. For imagery vividness, data is shown after removing effects of task performance. **C)** To assess the directionality of effects, we further obtained Granger causality (GC) (8-10 Hz) which correlated with imCOH across pairs of regions. Connections with conclusive evidence are shown in red and are further depicted in **D)** with their corresponding forward and backward GC estimates. **E)** imCOH values (8-10 Hz) are also depicted in circle plots. Error bars and shaded areas correspond to 95% HDI (A, B, D) or confidence (C) intervals. See **Fig S6, Fig. S7**, and supplementary data for full statistical report.

For directionality interpretation, we further estimated Granger causality (GC) between the same pairs of electrodes, yielding results highly consistent with imCOH (Fig. 4C). We observed modulations of bottom-up (STG→HPC, A1→VLPFC, STG→M1), top-down (DLPFC→MTG, STG→A1) and bidirectional connections (ACC↔STG, VLPFC↔HPC, parietal↔M1) (**Fig. S6**). By time-reversing one of the signals, we ruled out signal-to-noise directionality confounds (27) for connections originating in the STG toward HPC, A1, and M1, whereas for A1→VLPFC and DLPFC→MTG confounds remained a possibility (**Fig. S7**). This modulation profile is consistent with enhanced exchange of information between auditory regions and the rest of the brain during manipulation.

### 2.6. Alpha synchrony predicts imagery vividness and task performance

Next, we investigated whether the identified synchrony relates to behavior. We compared additional Bayesian mixed-effects models in which task performance, imagery vividness and relevant interaction terms were included as predictors on top of the previously tested block effects. We found evidence for interactions between block and task accuracy (β = 0.08, 95% HDI = [0.02 - 0.13]) and block and imagery vividness (β = −0.08, 95% HDI = [−0.13 - −0.03]) in the 8-10Hz range (**Fig. 4A**). With higher vividness, imCOH increased during recall and decreased during manipulation, whereas for better task performance it increased during manipulation and decreased during recall (Fig. 4B). GC analyses were again consistent with imCOH (**Fig. 4C**), revealing block/vividness and block/accuracy interactions in a distributed set of connections. For the vividness interaction, we ruled out signal-to-noise confounds in 8 out of 12 conclusively modulated pairs. In contrast, for task performance, we ruled out confounds in 4 out of 8 modulated pairs (**Fig. S7**). These results suggest that the relationship between brain synchrony and behavior depends on the context (manipulate vs. recall) and goals (accurate memory vs. vivid imagination) of the participant.

### 2.7. No evidence for manipulation effects on phase-amplitude cross-frequency coupling

The strength of higher-frequency neural activity often correlates with the phase of lower-frequency dynamics (28) (**Fig. 5A**). This phenomenon, known as phase-amplitude coupling (PAC), is thought to support working memory (10). To test whether PAC is linked to mental manipulation, we calculated the **modulation index (MI)** (29) as a measure of coupling between low-frequency phases (2-12 Hz in 2 Hz bins) and high-gamma amplitude for both baseline and delay separately.

**Fig 5.**
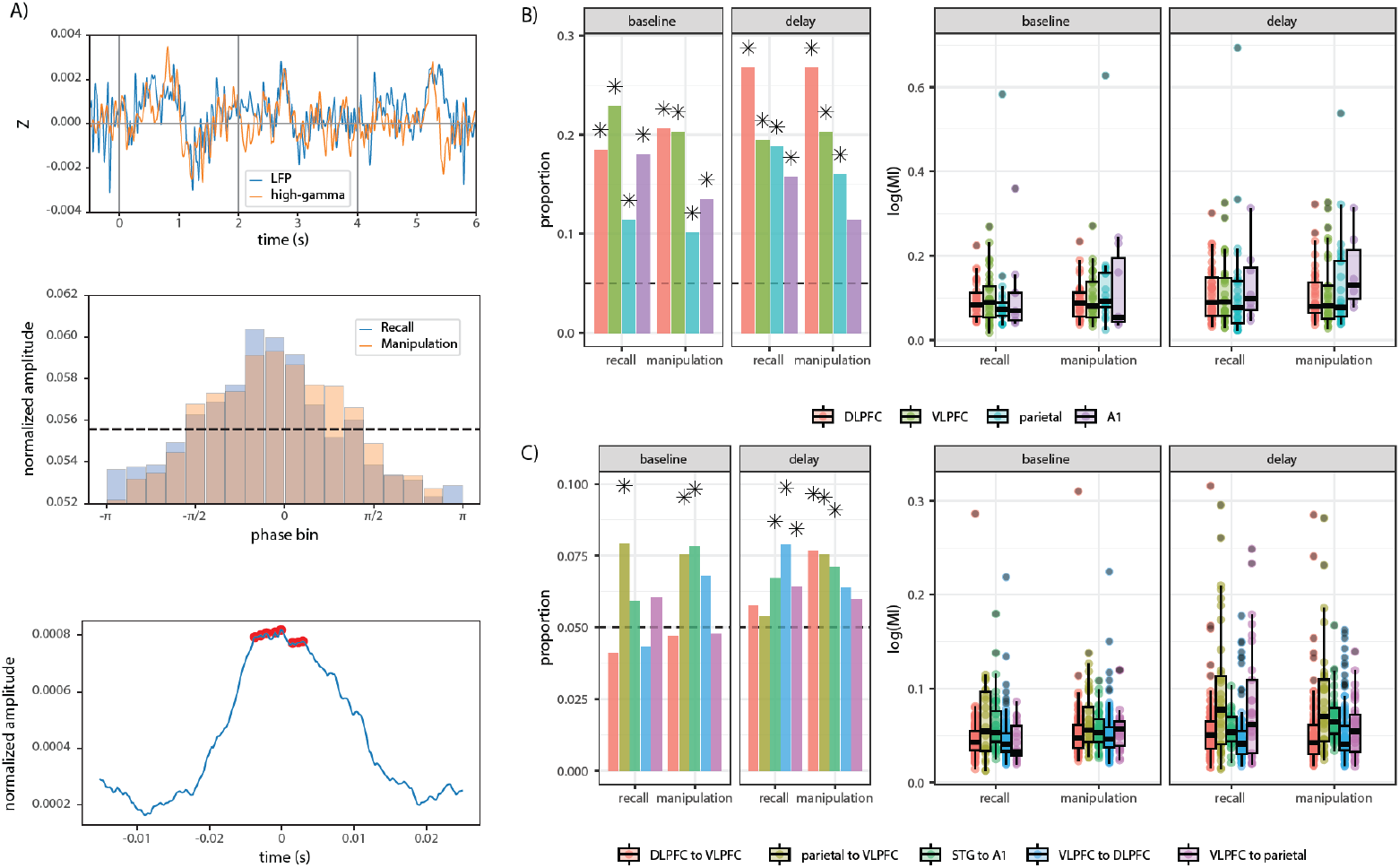
Phase-amplitude coupling. **A)** Example of a parietal channel with phase-amplitude coupling between low-theta and high-gamma. Coupling was observed in the correlation between local-field potentials (LFP) and high-gamma power in single trials (top), the distribution of high-gamma amplitude across phase bins (middle), and the high-frequency peaks (red dots) around the preferred phase of the mean low-frequency waveform (bottom). The dashed line marks the uniform amplitude distribution. **B)** Local and **C)** long-range PAC was present among a few frontoparietal and auditory ROIs and did not substantially change in its proportion (left) or strength (right) between recall and manipulation. Dashed lines indicate chance level for proportions of significant channels. See **Figures S8-11** and supplemental datasets for full statistical report.

We estimated **local PAC** within each electrode and **long-range PAC** between pairs of electrodes belonging to different ROIs. We identified electrodes or pairs with significant PAC by comparing observed values with surrogate null distributions. To avoid spurious effects of non-sinusoidal waveforms (30, 31), we identified true PAC as the cases where at least three high-frequency cycles could be observed in the average waveform at the preferred phase (32) (**Fig. 5A**). For statistical analysis, we used Bayesian mixed-effects models in which subject and region (or pair) were used as grouping variables for random intercepts and slopes (**table S6**). We included period (delay vs. baseline), condition (manipulation vs. recall), and their interaction as predictors. We compared models with and without these predictors and their corresponding random-effect terms (**table S6**). We ran two separate analyses: One for PAC strength (Gaussian) and another for the proportion of PAC channels (or pairs) per region (logistic).

We found local PAC in the dorsal stream (parietal, VLPFC, DLPFC) and the primary auditory cortex exclusively for low-theta (2-4 Hz) (**Fig. 5B & Fig S8**). Bayesian analyses revealed no conclusive effects of period (strength: β = −0.003, 95% HDI = [−0.21 - 0.21]; proportions: β = 0.10, 95% HDI = [−0.56 - 0.70]), block (strength: β = −0.002, 95% HDI = [−0.16 - 0.15]; proportions: β = −0.36, 95% HDI = [−1.22 - 0.31]) or an interaction between these variables (strength: β = −0.0005, 95% HDI = [−0.14 - 0.14]; proportions: β = −0.13, 95% HDI = [−0.87 - 0.58]).

Similarly, the proportion of pairs with long-range PAC was above chance for bidirectional parietal-VLPFC and DLPFC-VLPFC connections, and a unidirectional connection from STG (phase frequency) to A1 (amplitude frequency) (**Fig. 5C & S10**). Again, PAC was restricted to the low-theta frequency range. There was no evidence for effects of period (strength: β = −0.001, 95% HDI = [−0.16 - 0.16]; proportions: β = −0.11, 95% HDI = [−0.64 - 0.35]), block

(strength: β = −0.002, 95% HDI = [−0.12 - 0.12]; proportions: β = −0.11, 95% HDI = [−0.53 - 0.34]), or their interaction (strength: β = −0.005, 95% HDI = [−0.10 - 0.10]; proportions: β = - 0.10, 95% HDI = [−0.52 - 0.32]). Furthermore, there was no evidence for effects of vividness and task performance on channel proportions or PAC strength (**tables S7 & S8**).

## Discussion

With high spatiotemporal detail, we show that mental manipulation modulates neuronal dynamics in a wider frequency and anatomical range than previously suggested by non-invasive studies. We report two main effects: a) A shift in the balance between slower and faster neuronal dynamics in auditory areas, and b) behaviorally relevant modulations of brain-wide theta / alpha synchrony. These findings suggest an important role of local inhibition and neural synchrony in the coordination of sensory, memory, and control networks during manipulation.

Compared to baseline, power in sensory and dorsal areas increased in higher frequencies and decreased in lower frequencies. In contrast, power in ventral areas increased in lower frequencies and decreased in higher frequencies, resembling low-frequency modulations previously interpreted as gating (17). These frequency tradeoffs might reflect changes in neuronal excitability, where lower-frequency enhancements point to inhibition. This relationship is consistent with the negative correlation between beta and gamma bursts during WM in non-human primates (33); aligns with theoretical work assigning a “clearance” role to alpha and beta oscillations (34); and has been described with the slope of the (1/f) power spectral density taken to reflect excitation / inhibition balance (23). Consistent with this idea, manipulation effects on lower- and higher-frequency dynamics might index shifts in neuronal excitability. This is most evident in A1 and MTG, where manipulation enhanced lower frequencies and suppressed higher frequencies, likely reflecting inhibition of current sensory representation (34) to facilitate their transformation.

Importantly, we found that neuronal dynamics carried information about the context and content of WM. However, while melody identity was easily retrieved from encoding-period dynamics, it was difficult to decode from delay-period signals. This is likely due to the sharp drop in auditory cortex activity and suggests imagined sounds might be represented in fine-grained patterns of neuronal ensemble dynamics, possibly intermingled within small patches of cortex, which are difficult to detect with macroelectrodes. In a previous non-invasive study using the same task, we decoded melody identity from delay-period activity with above-chance accuracy (21). In that study, the number of participants (N=71) and trials (N=60) was larger, but accuracy was not significantly higher (∼55%). This highlights the challenge of detecting neuronal representations of mental objects with macroelectrodes and non-invasive techniques.

Consistent with our hypothesis, we found a narrow-band increase in high-theta / low-alpha (6-10 Hz) synchrony between auditory and non-auditory areas, likely reflecting enhanced exchange of control signals across the brain. Alpha oscillations are proposed to reflect pulsed inhibition (35) and seem ideal for network regulation: They are ubiquitous, increase with WM load (36), are enhanced during distractor suppression (11), and are prominent in auditory cortex, especially in secondary areas (37).

Intriguingly, alpha synchrony had a complex relationship with behavior. With higher reported vividness, synchrony increased during recall but decreased during manipulation, while the opposite was true for task performance. An explanation might be that, during manipulation, higher alpha synchrony pushes the network towards inhibition. However, for vivid imagination, auditory neurons need to actively engage, making decreases in synchrony helpful for vividness. Conversely, during recall, alpha synchrony and inhibition are reduced, which may make the network more susceptible to environmental noise. In this case, higher synchrony might help participants suppress interference and rely more on internal memory processes for successful imagery. Consequently, imagery vividness might rely on an optimal level of network synchrony that balances internal and external signals with the need for their maintenance or update.

Furthermore, note that imagery is not equivalent to WM and is only one of several possible maintenance strategies (22). In our study, participants primarily used imagery to perform the task, but they may have also relied on strategies like passive retention, at least in some trials. This would explain why the interaction between block and task performance was in the opposite direction compared to block and vividness. Thus, when imagery is not involved, higher network inhibition via alpha synchrony might help suppress conflicting sensory information during manipulation, leading to better task performance. This might be the mechanism by which parietal theta stimulation improves mental manipulation (13). These findings suggest imagery and short-term memory can impose conflicting excitability demands, which in turn depend on whether representations need to be held or updated. Thus, individuals who can optimally regulate network synchrony may benefit most from imagery in WM.

Another proposed mechanism for cognitive control is phase-amplitude coupling (PAC) (38), which we detected both within and among frontoparietal and auditory areas. However, the lack of manipulation effects indicates it may not be relevant for this function. This contrasts with previous reports of enhancement by WM load. In one study, the effect was observed in a minority of electrodes and was mainly driven by one participant (39). Another study used microelectrodes, suggesting PAC might be better observed at the level of microcircuits (10). Moreover, we controlled for spurious PAC arising from waveform shape, suggesting some of the reported effects (39, 40) might have been influenced by it. The extent of PAC involvement in WM control is yet to be determined.

To conclude, we provide evidence that neuronal synchrony and sensory inhibition play an important role in mental manipulation. These results provide an empirical ground for future experimental and theoretical work addressing how exactly neural dynamics aid cognitive control, how they interact with the neuronal ensembles that carry mental representations, and how they might be targeted in neurostimulation protocols for cognitive and neuropsychiatric disorders where WM and imagination are disrupted.

## Methods

### Participants

We included 30 patients undergoing presurgical epilepsy monitoring (N = 30, 16 female, age = 37.32 +/− 11.91 SD). We excluded patients with chance or below-chance task performance, low IQ, or low memory capacity. Patients gave informed written consent and were tested at different locations in the United States and Europe. Electrode coverage was tailored to each patient and determined solely by clinical needs. In 23 participants, we measured musical expertise with a short questionnaire asking for years of musical training as well as other questions not relevant for the current report (played musical style, etc.). The study was approved by each hospital’s internal review board, the University of California (Berkeley), and the Ethics Committee for the Capital Region of Denmark. The research was conducted according with the declaration of Helsinki.

### Stimuli

Stimuli consisted of short three-note melodies using piano sounds synthesized with MuseScore (v3.6.2) and normalized to peak amplitude. These sounds formed a major chord arpeggio (musical pitch: A3, C#5, E6; Fundamental frequency: 220Hz, 554Hz, 1318Hz) and were arranged in ascending order in melody 1 (A-C#-E) and descending order in melody 2 (E-C#-A), with a 500 ms inter-onset interval. We also included two foil test melodies: A-E-C# and E-A-C#. Sounds were presented through loudspeakers at a comfortable volume.

### Procedure

Participants completed a music background questionnaire before participating in the experiment. There were two types of setups. For 19 patients, we used the general-purpose BCI2000 software for synchronized auditory stimulus presentation, gaze tracking, behavioral feedback, and sEEG signal acquisition (41). All auditory stimuli were presented using a set of wired speakers (Logitech Multimedia Speakers Z200) positioned approximately 70 cm in front of the participants. Continuous monitoring of gaze fixation using a Tobii Pro Fusion Eye tracker sampling at 120 Hz (Tobii AB, Stockholm, Sweden) served as a secondary validation of task attendance. A Nihon Kohden JE-120A amplifier (Nihon Kohden Corp., Tokyo, Japan) sampled the SEEG and auditory onset (g.TRIGbox, g.tec, Austria) at 2000 Hz to ensure synchronization between auditory stimulus presentation and the resulting neural signals. The acquired data was stored in BCI2000 format for post-hoc analysis. For the remaining participants, the task was presented on a laptop with Psychopy software (42) using the laptop’s loudspeakers for stimulus delivery.

Before the onset of each trial, participants first saw the trial number (e.g., trial 32 / 48) for 2.5 seconds at the center of the screen (**Fig. 1A**). At trial onset, participants listened to either melody 1 or 2 while the word ‘listen’ appeared simultaneously at the center of the screen. At 2 s, the word ‘imagine’ appeared on the screen and participants were instructed to vividly imagine the previous melody in the **recall** condition, or its inverted version in the **manipulation** condition (i.e., melody 1: A-C#-E becomes melody 2: E-C#-A and vice versa). At 4 s, the word ‘listen’ appeared again on the screen and a test melody was presented which could be the same as the first one (50%), its backward version (25%), or a foil melody (25%). Participants had to press “1” on the keyboard if the melody was the same as the first one (recall block) or its inversion (manipulation block), or press “2” otherwise. The two conditions (recall vs. manipulation) were presented in separate blocks with 48 trials each. For two participants, 60 trials per condition were used instead. A practice session was included at the beginning of each block with 2 trials each. At the end of each block, participants were asked to rate on the computer from 1 (not vivid at all) to 5 (extremely vivid) how vivid their imagination was. For all except two participants, a localizer block (∼5 min, not analyzed here) was presented before the main task in which the three sounds were played in pseudorandom order. An additional auditory recognition memory task (∼5 min, not analyzed here) was included either before the localizer or after the main task. Block order was counterbalanced across participants. The whole experiment lasted ∼30 min.

### Behavioral analyses

All statistical analyses were done in R (43) unless stated otherwise. We used mixed-effects logistic regression (“lme4” package, 44) to estimate the rate of correct responses and the effect of block (recall vs. manipulation), vividness, and musical expertise (musicians vs. non-musicians) on them (**table S1**). We defined musicians as participants with 3 or more years of musical training. For analysis of vividness ratings, we used ordinal logistic regression in the form of cumulative-link mixed-effects models (“ordinal” package, 45) and included block, musical expertise and corresponding interactions as fixed-effects predictors (**table S2**). We entered subject as grouping variable for random intercepts and slopes in all models. We used likelihood ratio tests to compare models with and without a predictor to test its effect on the dependent variable. We employed an incremental approach starting with an intercept-only model and ending with a full model with all predictors and interactions. We obtained 95% confidence intervals for parameter estimates using the Wald method (“broom mixed” package 46) and predicted marginal means and their 95% confidence intervals using covariance matrices as implemented in the “effects” package (47).

### iEEG preprocessing

iEEG analyses were primarily performed using MNE-Python (48). Data were exclusively recorded from stereotactic EEG electrodes. After converting the data to BIDS format, we visually inspected the recordings and excluded channels with epileptic activity (inter-ictal spikes, slowing), corrupted data, or located outside of the brain. We applied bipolar re-referencing to neighboring electrodes within each electrode shaft by subtracting the outer from the inner contacts, resulting in one less channel per shaft. We obtained electrode locations based on post-operative computerized tomography images corregistered with individual MRI scans (49). The coordinates were translated into standard MNI space for visualization. Bipolar pairs with both channels located in white matter were excluded from further analyses. To account for bipolar re-referencing, the final location of a contact was determined as the mid-point between its original location and that of its neighbor. After resampling (600 Hz) and power-line noise removal (60 Hz USA, 50 Hz Europe), we epoched the data from -2 to 8 s relative to trial onset.

The anatomical location of each contact was visually determined and verified with the aid of an expert neurologist. We selected 11 regions of interest (ROI) for analysis as follows: A1 (Heschl’s gyri / transverse temporal gyri), STG (anterior and posterior superior temporal gyri), STS (superior temporal sulcus), MTG (anterior and posterior middle temporal gyrus), HPC (anterior and posterior hippocampus), PHC (parahippocampal), parietal (superior parietal lobe, inferior parietal lobe, angular gyrus, supramarginal gyrus, temporo-parietal junction, intraparietal sulcus), VLPFC (inferior frontal gyrus, frontal operculum, inferior frontal sulcus), DLPFC (middle frontal gyrus, superior frontal sulcus, frontal eye field), M1 (pre-central gyrus) and ACC (ventral and dorsal portions of the anterior cingulate) (**Fig. 1B & Fig. S1**).

### Time-frequency analyses

We estimated time-frequency representations using Morlet wavelet decomposition with 36 center frequencies spanning seven canonical frequency bands: Low-theta (1-3 Hz – 0.5 Hz bins), high-theta (4-7 Hz – 1 Hz bins), alpha (8-12 Hz – 1 Hz bins), low-beta (13-19 Hz – 2 Hz bins), high-beta (21-29 Hz – 2Hz bins), low-gamma (31-55 Hz – 5 Hz bins), and high-gamma (70-160 Hz – 10Hz bins). For each subject, channel, and center frequency, we log-transformed power values and z-scored them by subtracting the mean of the baseline (−1.5 s to −0.5 s) across all trials and dividing by the corresponding standard deviation. Subsequently, we further subtracted the baseline of each trial, making sure power modulations reflected task-related activity. We decimated the power time courses (25 ms steps) and smoothed them (100 ms time windows) for further analysis.

We averaged the resulting values across center frequencies and trials, resulting in one time course per canonical band, channel, and condition. These values were used for statistical analyses as described next. For each ROI, time point, and frequency band, we fitted Bayesian mixed-effects models (“MCMCglmm” package, 50) with subject as a grouping variable for random intercepts and slopes (**table S3**). For estimation, we used Markov Chain Montecarlo (MCMC) with 20.000 samples (1.000 burn-in). To assess model convergence, we used Lilliefors’ tests to determine whether Markov chains were possibly deviating from normality, which we then inspected visually. All reported models converged. Priors for fixed-effects parameters were set to MCMCglmm default values, centered at 0 with an uninformative variance of 1000. The structure of random-effects covariances was set to the same value for all terms, with priors consisting of an inverse-Wishart distribution centered at 1 with dispersion parameter 0.2. This choice of priors ensured the convergence of all models.

We fitted three different models. An intercept-only model, a model with block (manipulation vs. recall) as a fixed-effect predictor, and a model with random slopes (**table S3**). We compared the models by calculating Deviance Information Criteria (DIC) and deriving model weights (or probabilities) from them (51). We obtained 95% highest density intervals (HDI) from the posteriors and estimated model predictions and their 95% HDI intervals with the “bayestestR” package (52). We regarded a block effect as conclusive if the HDI of the block coefficient excluded 0 (in the model with random slopes) and the cumulative probability of models including block was 0.95 or higher.

### Decoding

We used logistic regression to decode task variables from band-power estimates. For each subject, we first used all the ROIs (global decoding) to predict block (manipulate vs recall) or melody identity (mel1 vs. mel2). We then repeated the same analyses separately for each ROI (local decoding). As features or predictors in the models, we entered each time point at each channel to take advantage of temporal patterns. All analyses were done for the encoding (0 - 2 s) and delay period (2 – 4 s) and for each canonical frequency band separately. In addition, melody decoding was done individually for each block. We used balanced accuracy as scoring metric in a five-fold cross-validation procedure where we took the average of the five folds as output. To reduce variability in accuracy outcomes, we repeated the cross-validation procedure 10 times with different random seeds and took the mean of the repetitions.

For statistics, we centered accuracy scores by subtracting the chance level (0.5) and fitted Bayesian mixed models as described before. We fitted intercept-only models to test whether accuracies were different from chance, including subject and frequency band as grouping variables for random terms (**table S4**). We inspected random effect coefficients to characterize decoding accuracies for different bands and ROIs. Assessment of conclusive evidence, model convergence, model comparisons, and estimation of HDI were done as described before.

### Connectivity analyses

Using the “MNE_connectivity” library in Python (53), we estimated directed synchrony between pairs of channels from different ROIs within the same subject as the imaginary part of coherence (imCOH)

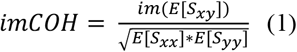

where *E*[*S*_*xy*_], *E*[*S*_*xx*_], *E*[*S*_*yy*_] are the expected values across trials of cross and power spectral densities for a pair of channels *x* and y.

We obtained estimates for the baseline (−2 - 0s) and the delay period (2 - 4s) and for each condition (manipulation vs. recall) separately, in a frequency range from 2 to 30 Hz in 2Hz bins. Due to the ambiguous interpretation of the sign of the output, we took the absolute value of the estimates (|imCOH|) for further analysis. To determine statistical significance for a given channel pair, frequency bin, and time period, we compared the result with a surrogate null distribution obtained by randomly shuffling trial labels of one of the signals 200 times. We deemed a connectivity value as significant if it was equal to or higher than the 95% percentile of the surrogate distribution.

For statistical analyses, we selected pairs with significant values in any of the conditions or time periods. We log-transformed the values, replacing the few cases with negative infinity (i.e., log(0)) by the next lowest value in the data. We z-scored these outcomes and computed the difference between delay and baseline estimates. This allowed us to account for baseline levels of synchrony, singling out connectivity effects specific to the delay period (26).

For statistical analysis, we used Bayesian mixed-effects models including subject and connection (i.e., pair of ROIs) as grouping variables for random intercepts and slopes. We fitted one model for each frequency bin. As in the case of power analyses, we used DIC and model weights for comparisons. We fitted several models incrementally, starting with an intercept-only model and adding block, task performance (i.e., percent correct), imagery vividness, and corresponding interactions, until we reached the full model (**table S5**). We obtained HDI and model predictions as described above. We deemed evidence for a given predictor as conclusive if its 95% HDI excluded zero (in the model that first introduced the predictor and its random slopes), and the cumulative probability of models testing the predictor was 0.95 or higher (when compared with the adjacent null model that excluded the term, see **table S5** for details). Model convergence was ensured as described above.

To estimate the connections modulated by task variables, we inspected the posterior distribution of random effect coefficients, as obtained from MCMC, after adding corresponding fixed effect coefficients. This approach optimally handles the multiple comparison problem across connections due to the regularizing properties of Bayesian mixed-effects models (54). We deemed evidence for modulation of a specific connection as conclusive if its 95% HDI excluded 0.

To gain a better grasp of the directionality of interactions, we estimated Granger Causality between the same channel pairs following the MNE_connectivity multivariate implementation based on

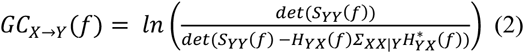

Where X is a set of seed signals, Y is a set of target signals, *f* is a given frequency, *Σ* is the residuals’ covariance from an autoregressive model, *H* is a spectral transfer function linking the residuals with the cross-spectral density between the signals, and S corresponds to *Σ* transformed by *H*. Note that, although this implementation is multivariate, we used it to compute GC between pairs of signals only.

GC estimates were subject to the same statistical procedures as imCOH with the only difference that connections were broken into forward and backward directions. We evaluated the consistency of GC estimates by computing the mean of forward and backward values for each connection, as obtained from random effects coefficients, and correlating the output with imCOH estimates. These checks revealed high consistency between GC and imCOH (**Fig. 4C**).

Differences in GC strength between the forward and backward direction for a pair of signals might reflect true asymmetries or artifacts stemming from differences in signal-to-noise ratio. To diagnose the presence of artifactual directionality, we estimated time-reversed GC and repeated the analysis (27). We regard connections with true directionality effects as those in which the difference between forward and backward estimates changes direction after time reversal (**Fig. S7**).

### Cross-frequency coupling

We calculated phase-amplitude cross-frequency coupling (PAC) between a set of lower frequencies (2-12 Hz in 2 Hz bins) and broadband high-gamma power (before decimation and smoothing). We used the Modulation Index (MI) (29) as a PAC metric, as follows:

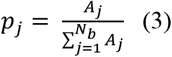

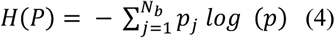

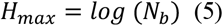

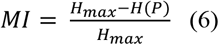

Where *A*_*j*_ is the high-frequency amplitude corresponding to a particular low-frequency phase bin *j, N*_*b*_ is the number of frequency bins, *p*_*j*_ is the amplitude normalized to the interval [0,1], *H(P)* is the entropy of the amplitude distribution across phase bins, and *H*_*max*_ is the entropy of a uniform amplitude distribution. Thus, MI becomes larger as the amplitude distribution deviates from uniformity and concentrates at certain phases. Using 18 bins (π / 9 radians each), we estimated MI for baseline and delay, and recall and manipulation separately, after concatenating the phases of all trials for a specific period and condition. We estimated both local PAC between phases and amplitudes of the same channel, and long-range PAC between phases and amplitudes of channels in different ROIs within the same subject. To determine statistical significance, we compared the resulting values with surrogate distributions obtained by shuffling trial labels of the amplitude signal 200 times. An estimate was deemed significant if it was equal to or larger than the 95% percentile of the surrogate distribution. The resulting MI estimates were then log-transformed (after adding 1 to avoid infinite values: log(MI + 1)) and z-scored.

Since spurious PAC might arise due to non-sinusoidal waveform shapes (30, 31), we used the procedure in (32) to identify true PAC. Briefly, we identified the preferred phase bin for each channel or pair (i.e., the bin with the largest amplitude). Then, we took all the instances of that bin and found the peak of the high-amplitude signal within each instance. Subsequently, we selected time windows centered at these amplitude peaks, corresponding to one cycle of the current phase frequency, and took the average across them. In the resulting mean waveform, we identified time-domain peaks around the preferred phase within a time window corresponding to plus or minus 3 cycles of the minimum amplitude frequency (70 Hz) (**Fig. 5A**). We defined true PAC as cases where at least three such peaks could be identified.

For statistical analyses, we first used a binomial test with a significance level of 0.05 to determine the ROIs (or ROI pairs) with a proportion of significant channels higher than expected by chance. We then selected significant ROIs (or ROI pairs) for further analysis. We tested for changes in the proportion of significant channels as a function of period (baseline vs. delay), block (manipulation vs. recall), and their interaction using Bayesian mixed-effects logistic regression including ROI (or ROI pair) and subject as grouping variables for random intercepts and slopes (**table S6**). To assess possible associations with task performance and imagery vividness, we estimated a set of additional models for each period separately, starting with intercept-only models and incrementally adding block and the other predictors, as well as their interactions (**tables S7 & S8**). We followed the same testing approach for MI strength. In this case, however, we used Bayesian linear mixed models after selecting channels with significant MI. Assessment of conclusive evidence, model convergence, model comparisons, and estimation of HDI were done as described for connectivity analyses.

## Supporting information

data1 - band power statistics

data2 - connectivity statistics

data3 - PAC statistics

## Funding

This work was supported by NIH/NINDS R01-NS021135 (RTK); Brain Initiative NIH/NINDS U19-NS107609 (RTK), U01-NS108916 (PB) and U24-NS109103 (PB); CONTE Center NIH/NIMH P50-MH109429 (RTK); NIH/NIMH R01-MH120194 (JTW); NIH/NIBIB P41-EB018783 (PB) and R01-EB026439 (PB); the Independent Research fund Denmark (DQM); the Carlsberg Foundation (CF23-1491) (DQM) and (CF20-0239) (LB); Lundbeck Foundation (Talent Prize 2022) (LB); Linacre College of the University of Oxford (Lucy Halsall fund) (LB); Nordic Mensa Fund (LB); the Danish National Research Foundation (DNRF 117) (GFR, LB, PV); Fondazione Neurone (PB); and the Mutua Madrileña Foundation (GFR); The Research Council of Norway project 314925 (AOB) and RITMO Center of Excellence 262762 (AOB, AKS, TE). The funders had no role in study design, data collection and analysis, decision to publish, or preparation of the manuscript.

## Author contributions (CRediT)

DQM: Conceptualization, data curation, formal analysis, funding acquisition, investigation, methodology, project administration, software, validation, visualization, writing – original draft. TH: formal analysis, methodology, software, writing – review and editing. GFR: Visualization, writing – review and editing; LB: Conceptualization, writing – review and editing. AB: Data curation, investigation, writing – review and editing. AKS: Data curation, investigation, writing – review and editing. TE: Data curation, investigation. TRM: Data curation, investigation. MF: Data curation, investigation. OKM: Data curation, investigation. JTW: Data curation, investigation. PB: Data curation, investigation. MD: Data curation, investigation. JL: Data curation, investigation. RTK: Funding acquisition, project administration, data curation, resources, supervision.

## Data and code availability

Data and code will be shared upon acceptance in a peer reviewed journal.

## Conflict of interest

The authors report no conflict of interest.

## Notes

### Competing Interest Statement

The authors have declared no competing interest.

