## Supplemental Figures and Tables for "Brain dynamics of mental manipulation"

Supplemental Materials

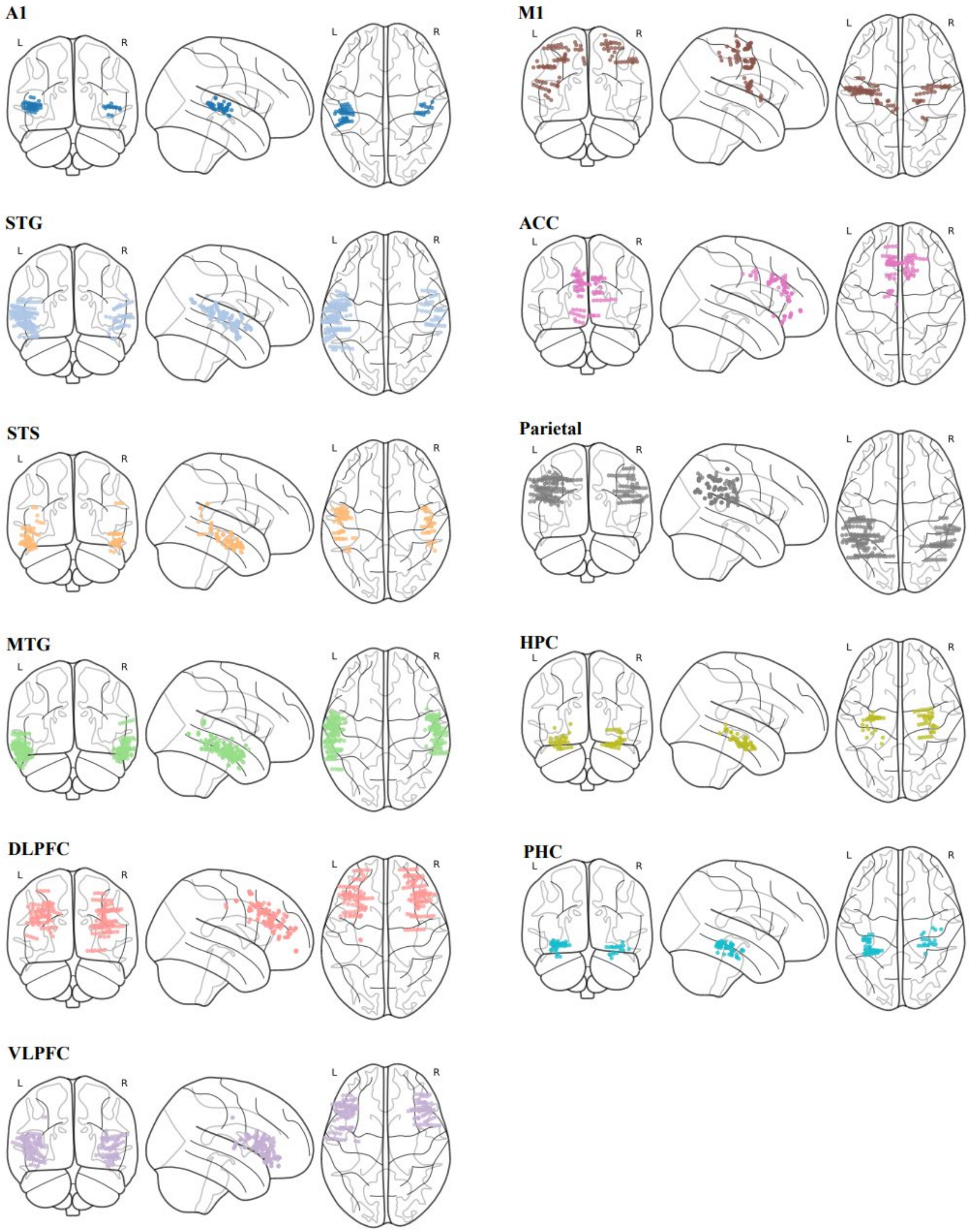


**Figure S1.** Electrode locations (normalized MNI space).

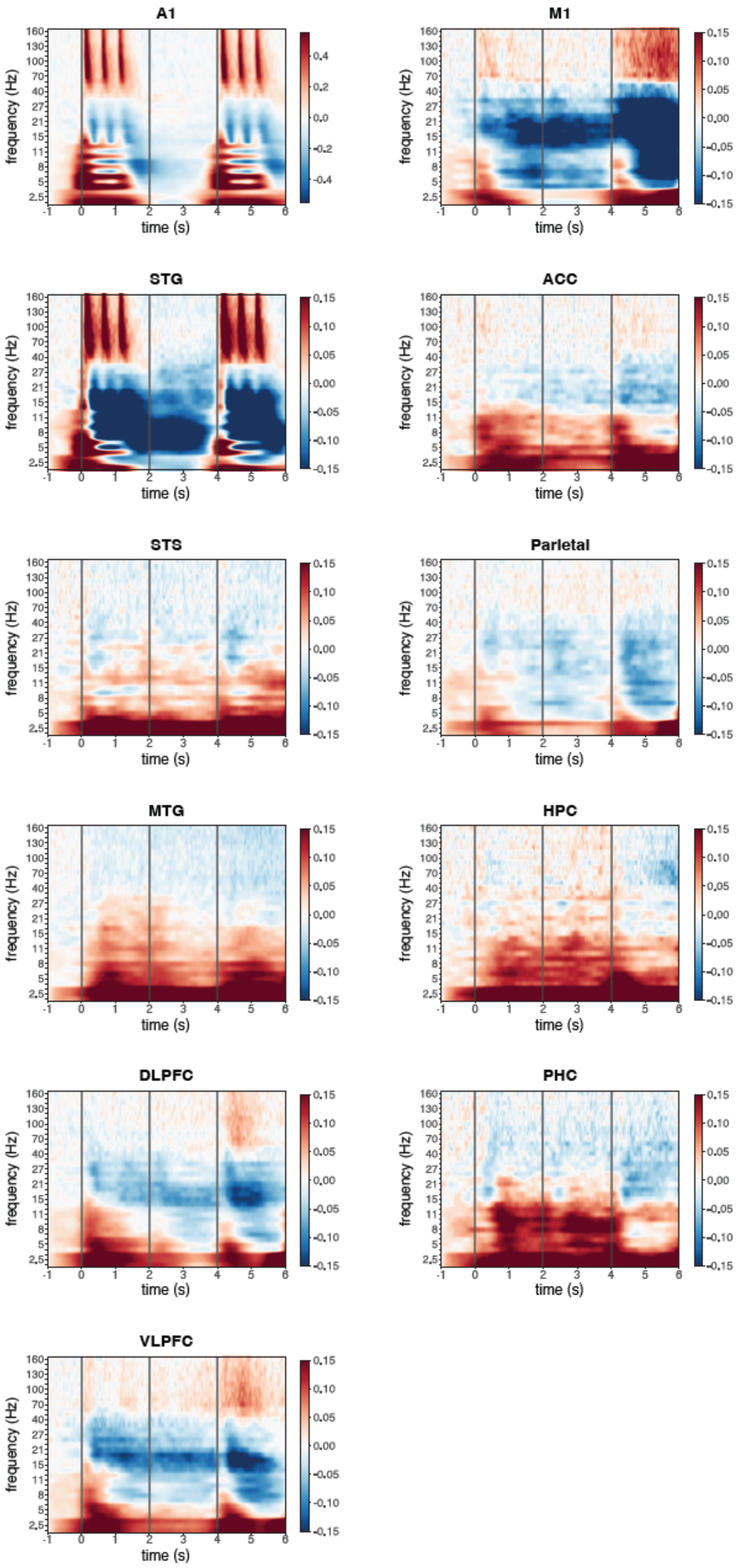


**Figure S2.** Average time-frequency representation of task-related activity. Vertical lines mark the onset of the first melody (0 s), the delay period (2 s), and the test melody (4 s).

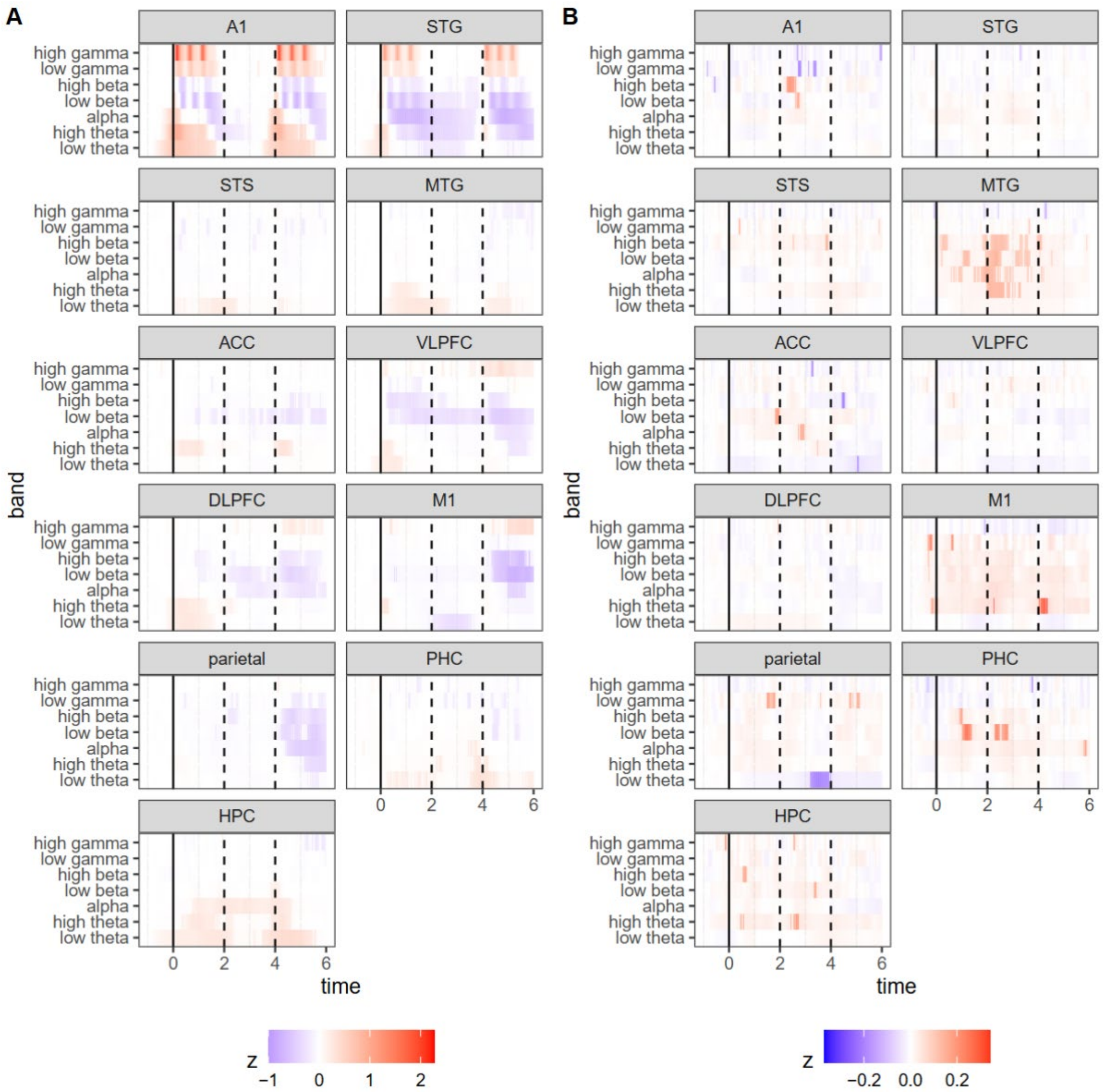


**Figure S3.** Band power coefficient estimates for **A)** the intercept and **B)** the block effect. Highlighted segments indicate 95% Highest Density Intervals (HDI) excluding 0. Dashed vertical lines mark the onset of the first melody (0 s), the delay period (2 s) and the test melody (4 s).

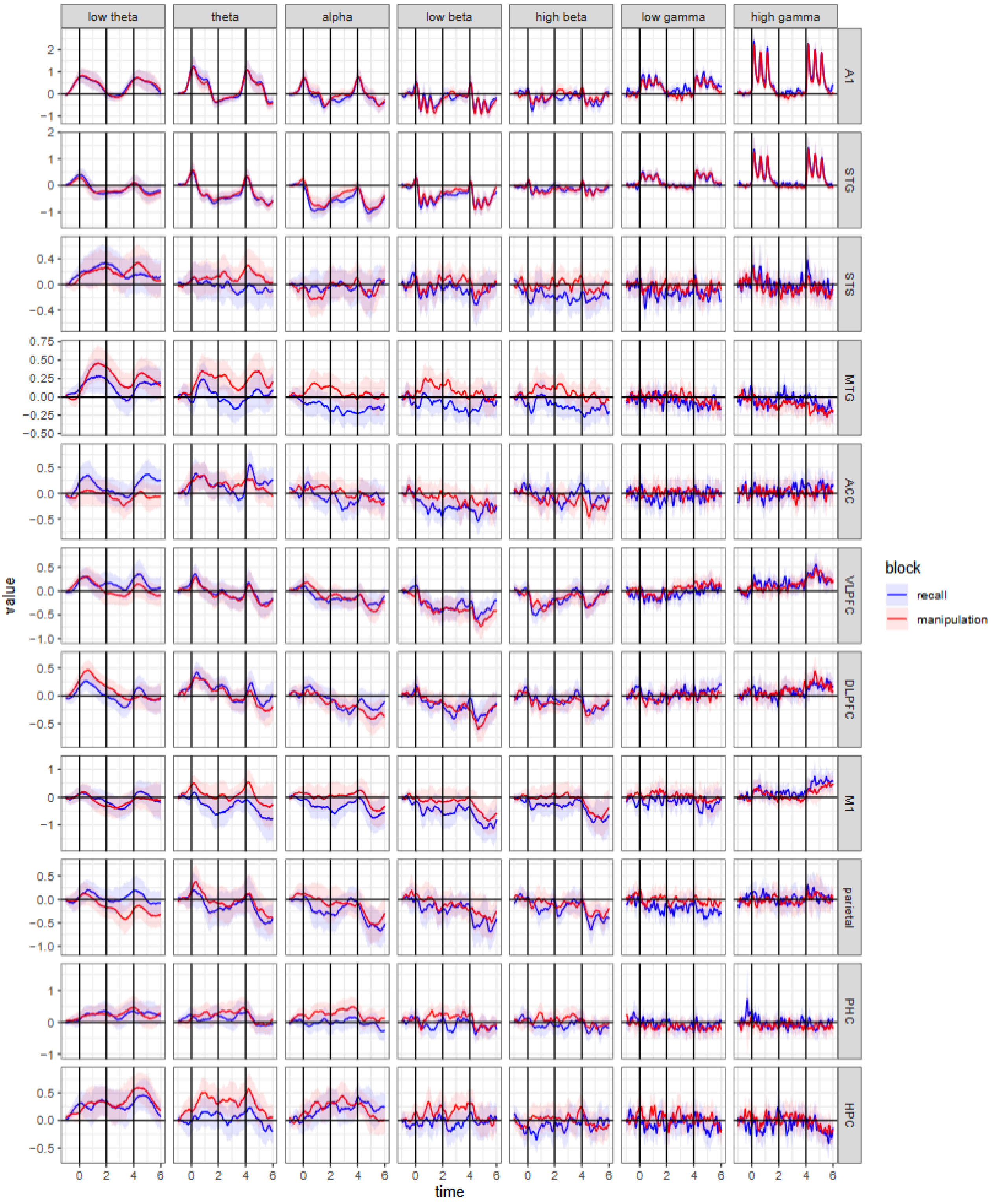


**Figure S4**. Band-power predictions obtained from the full model. Shaded areas depict 95% highest density interval (HDI). Vertical lines mark the onset of the first melody (0 s), the delay period (2 s), and the test melody (4 s).

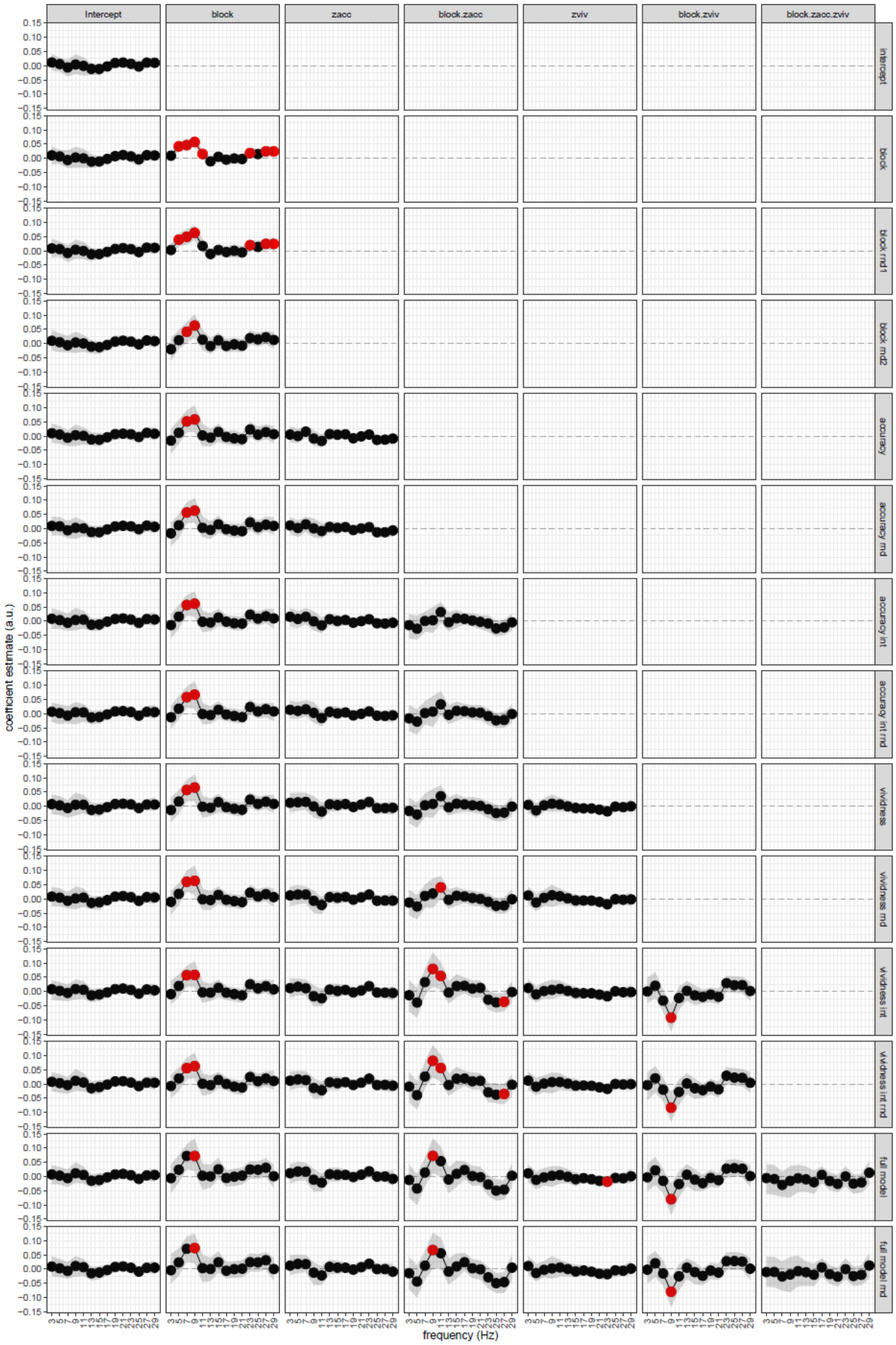


**Figure S5.** Imaginary coherence connectivity coefficients for different predictors (columns) across different statistical models (rows). Red dots indicate conclusive evidence and shaded areas 95% HDI. Zacc = z-scored accuracy, zviv = z-scored vividness, rnd = random effects, int = interaction. See table S2 for model structure. Full statistical report is provided as a separate dataset.

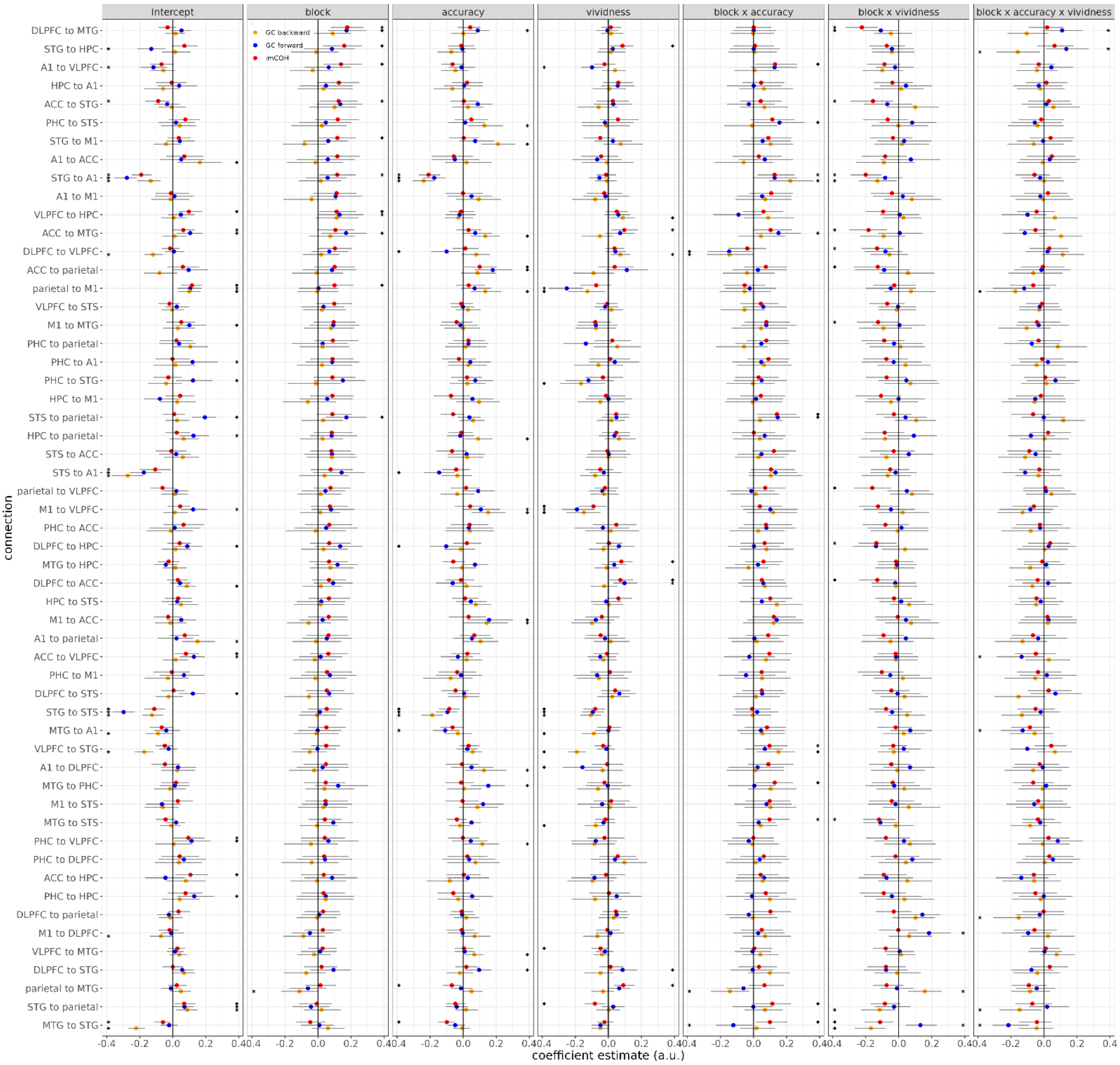


**Figure S6.** Random-effect coefficients from a full model with all main effects and interactions for the peak frequency bin (8-10 Hz), sorted by the strength of the block effect for imaginary coherence. Error bars represent 95% HDI. Asterisks mark 95% HDI excluding 0. Full statistical report is provided as a separate dataset.

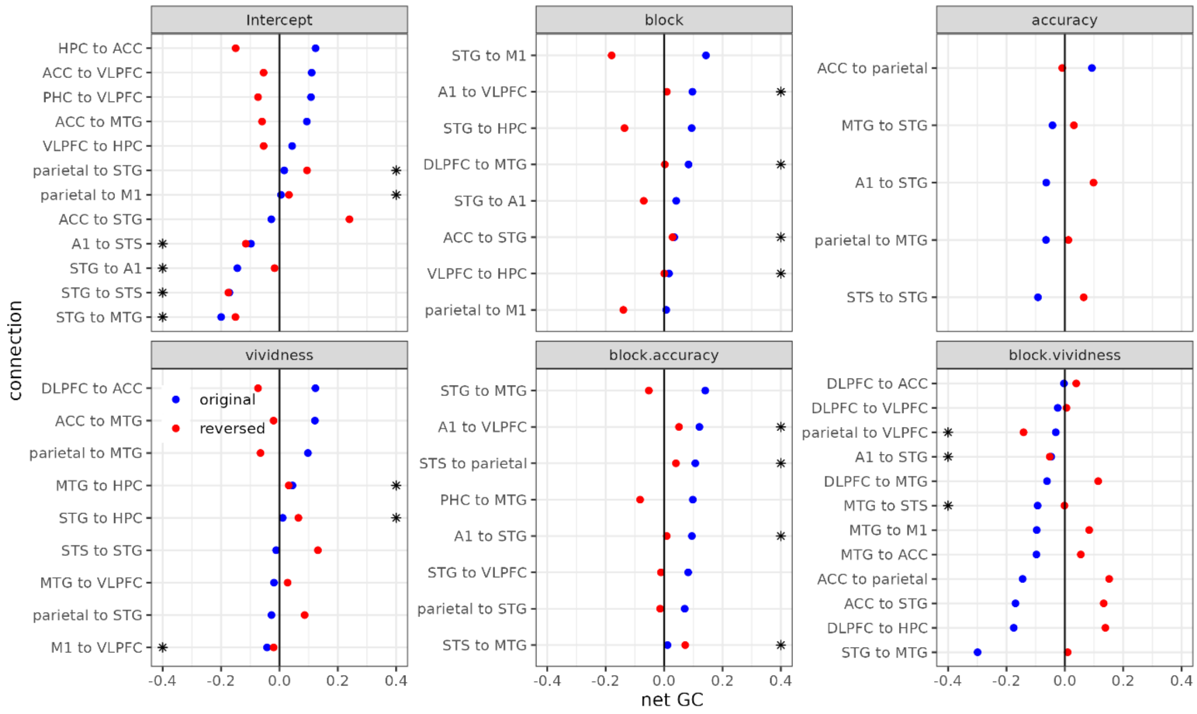


**Figure S7.** Directionality confound analysis comparing original with time-reversed Granger causality coefficients for pairs with conclusive imaginary coherence modulations. If true asymmetric directional interactions are present among two regions, time reversing one of the signals should reverse the direction of the effects. Conversely, if asymmetries are confounded by differences in signal-to-noise ratio between the regions, no reversal should occur. Asterisks mark region pairs with no directionality reversal.

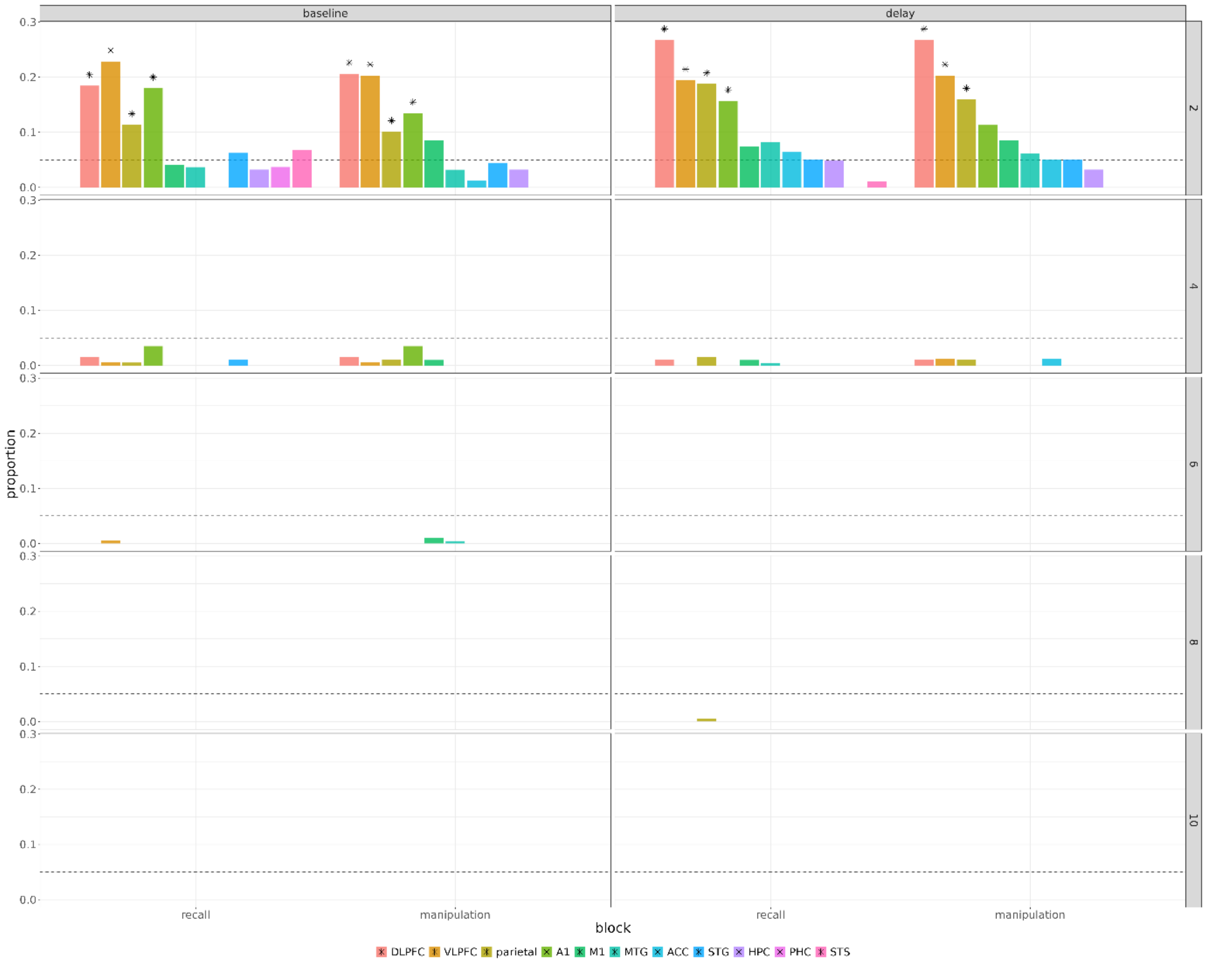


**Figure S8.** Proportion of significant local PAC channels for the two periods and different frequency bins (rows). Asterisks mark proportions significantly different from error level (dashed line, 0.05).

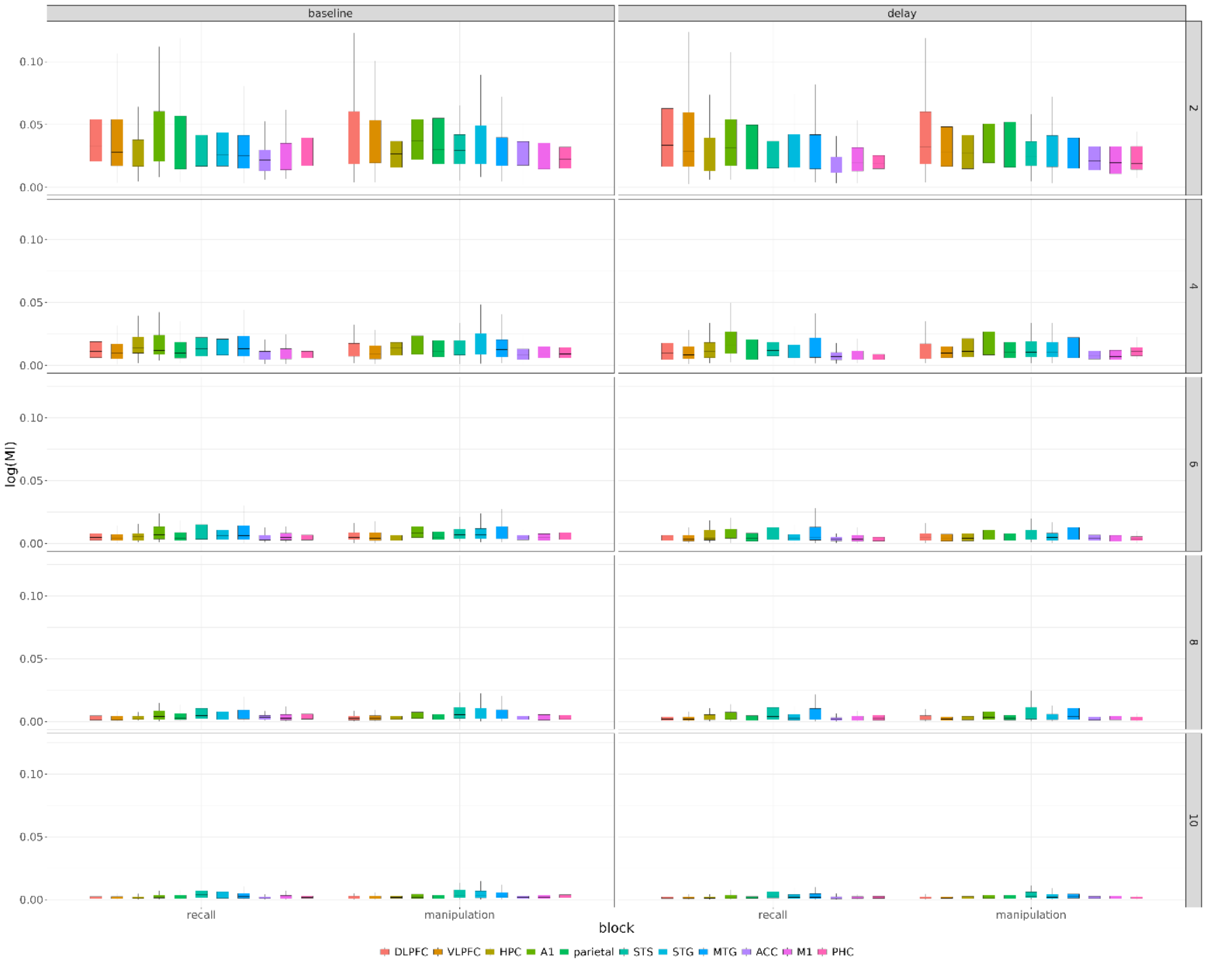


**Figure S9.** Boxplots (MDN +/- IQR) of local PAC strength (log modulation index) for the two periods and different frequency bins (rows).

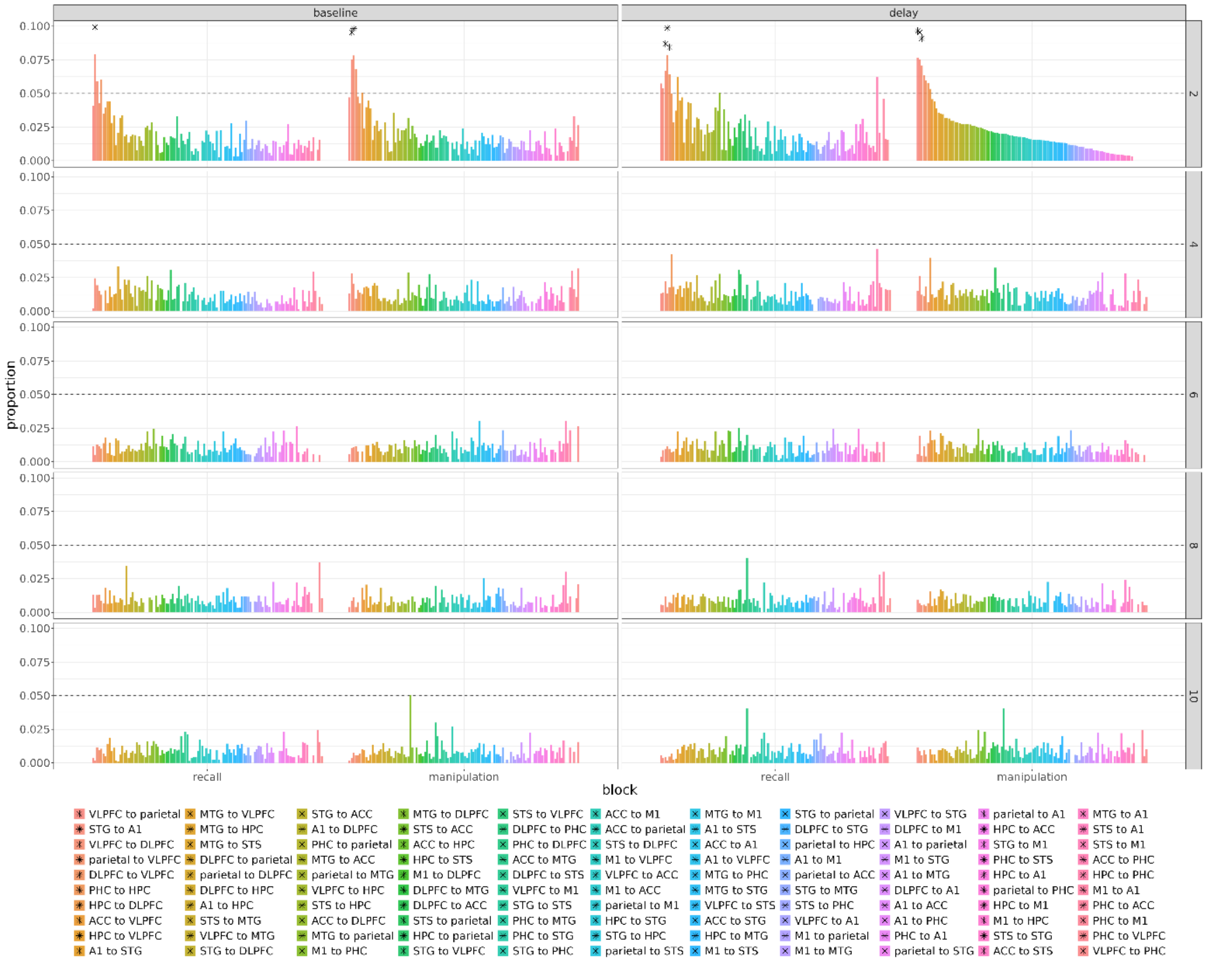


**Figure S10.** Proportions of significant long-range PAC channels for the two periods and different frequency bins (rows). Asterisks mark proportions significantly different from error level (dashed line, 0.05).

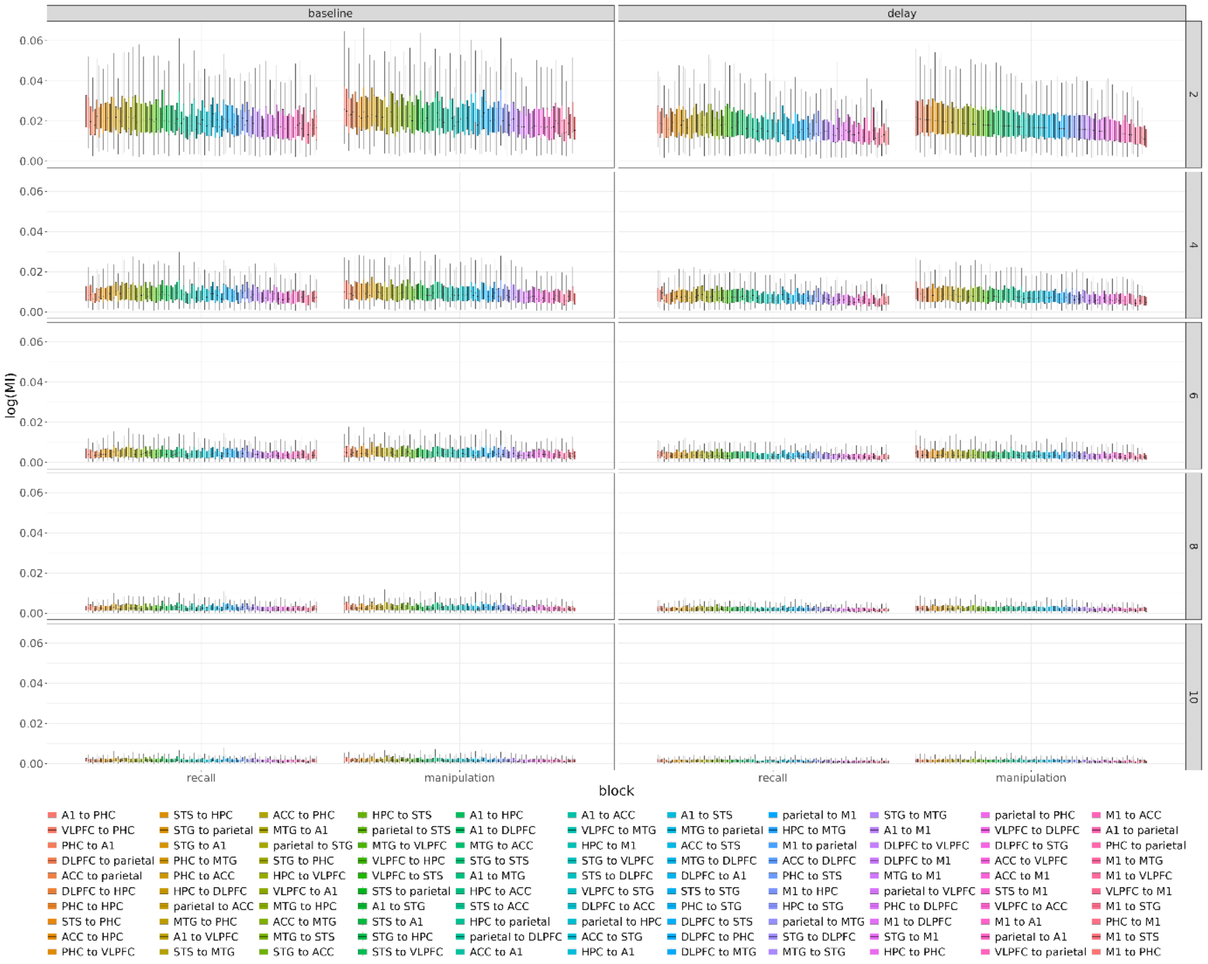


**Figure S11.** Boxplots (MDN +/- IQR) of long-range PAC strength (log modulation index) for the two periods and different frequency bins (rows)

| Behavioral accuracy (binomial) | | | | | | |
|---|---|---|---|---|---|---|
| **model** | **Null** | **formula** | $\chi^2$ | **Df (diff)** | ***p*** | ***AIC*** |
| m0 | - | accuracy ~ 1 + (1 \| subject) | - | - | - | 1517.3 |
| m1 | m0 | accuracy ~ 1 + block + (1 \| subject) | 68.39 | 1 | **< 0.001** | 1450.9 |
| m1r | m1 | accuracy ~ 1 + block + (1 + block \| subject) | 1.07 | 2 | 0.585 | 1453.8 |
| m2 | m1r | accuracy ~ 1 + block + vividness + (1 + block \| subject) | 7.72 | 1 | **0.005** | 1448.1 |
| m3 | m2 | accuracy ~ 1 + block + vividness + block:vividness + (1 + block \| subject) | 0.63 | 1 | 0.426 | 1449.5 |
| m4 | m3 | accuracy ~ 1 + block + vividness + block:vividness + expertise + (1 + block \| subject) | 3.08 | 1 | 0.079 | 1050.3 |
| m5 | m4 | accuracy ~ 1 + block + vividness + block:vividness + expertise + block:expertise + (1 + block \| subject) | 0.54 | 1 | 0.463 | 1051.7 |
| m6 | m5 | accuracy ~ 1 + block + vividness + block:vividness + expertise + block:expertise + vividness:expertise + (1 + block \| subject) | 2.08 | 1 | 0.149 | 1051.6 |
| m7 | m6 | accuracy ~ 1 + block + vividness + block:vividness + expertise + block:expertise + vividness:expertise + block:vividness:expertise + (1 + block \| subject) | 2.5 | 1 | 0.114 | 1051.1 |

**Table S1.** Statistical testing of behavioral accuracy. Each model was compared with a null model through likelihood ratio test. Models m4 to m7 were fitted with a reduced dataset because musical expertise information was not available for three participants, making AIC not directly comparable with models m0 to m3. Significant effects are highlighted in bold.

| Vividness ratings (ordinal) | | | | | | |
|---|---|---|---|---|---|---|
| **model** | **Null** | **formula** | $\chi^2$ | **df** | ***p*** | ***AIC*** |
| m0 | - | vividness ~ 1 + (1 \| subject) | - | - | - | |
| m1 | m0 | vividness ~ 1 + block + (1 \| subject) | 0.43 | 1 | 0.514 | 156.19 |
| m2 | m1 | vividness ~ 1 + block + expertise + (1 \| subject) | 1.42 | 1 | 0.23 | 122.2 |
| m3 | m2 | vividness ~ 1 + block + expertise + block:expertise + (1 \| subject) | 0.29 | 1 | 0.459 | 124.91 |

**Table S2.** Statistical testing of vividness ratings. Each model was compared with a null model through likelihood ratio tests. Models m2 to m3 were fitted with a reduced dataset because musical expertise information was not available for three participants, making AIC not directly comparable with models m0 and m1.

| Band power (Gaussian) | | |
|---|---|---|
| **model** | **Null** | **formula** |
| m0 | - | power ~ 1 + (1 \| subject) |
| m1 | m0 | power ~ 1 + block + (1 \| subject) |
| m1r | m1 | power ~ 1 + block + (1 + block \| subject) |

**Table S3.** Model structures for band power statistical comparisons. Full statistical report is provided as a separate dataset.

| Decoding accuracy relative to chance (Gaussian) | | |
|---|---|---|
| **model** | **Null** | **formula** |
| m0 | - | decoding accuracy ~ 1 + (1 \| subject) + (1 \| band) |

**Table S4.** Model structure for estimates of decoding accuracies. No model comparisons were performed.

| Connectivity (Gaussian) | | |
|---|---|---|
| **model** | **null** | **formula** |
| intercept | - | connectivity ~ 1 + (1 \| subject) + (1 \| connection) |
| block | intercept | connectivity ~ 1 + block + (1 \| subject) + (1 \| connection) |
| block rnd1 | intercept | connectivity ~ 1 + block + (1 \| subject) + (1 + block \| connection) |
| block rnd2 | intercept | connectivity ~ 1 + block + (1 + block\| subject) + (1 + block \| connection) |
| accuracy | block rnd2 | connectivity ~ 1 + block + accuracy + (1 + block\| subject) + (1 + block\| connection) |
| accuracy rnd | block rnd2 | connectivity ~ 1 + block + accuracy + (1 + block\| subject) + (1 + block + accuracy \| connection) |
| accuracy int | accuracy rnd | connectivity ~ 1 + block + accuracy + block:accuracy + (1 + block \| subject) + (1 + block + accuracy \| connection) |
| accuracy int | accuracy rnd | connectivity ~ 1 + block + accuracy + block:accuracy + (1 + block\| subject) + (1 + block + accuracy + block:accuracy \| connection) |
| vividness | accuracy int | connectivity ~ 1 + block + accuracy + block:accuracy + vividness + (1 + block \| subject) + (1 + block + accuracy + block:accuracy \| connection) |
| vividness rnd | accuracy int | connectivity ~ 1 + block + accuracy + block:accuracy + vividness + (1 + block \| subject) + (1 + block + accuracy + block:accuracy + vividness \| connection) |
| vividness int | vividness rnd | connectivity ~ 1 + block + accuracy + block:accuracy + vividness + block:vividness + (1 + block \| subject) + (1 + block + accuracy + block:accuracy + vividness \| connection) |
| vividness int rnd | vividness rnd | connectivity ~ 1 + block + accuracy + block:accuracy + vividness + block:vividness + (1 + block \| subject) + (1 + block + accuracy + block:accuracy + vividness + block:vividness \| connection) |
| full model | vividness int rnd | connectivity ~ 1 + block + accuracy + block:accuracy + vividness + block:vividness + accuracy:vividness + block:accuracy:vividness + (1 + block \| subject) + (1 + block + accuracy + block:accuracy + vividness + block:vividness \| connection) |
| full model rnd | vividness int rnd | connectivity ~ 1 + block + accuracy + block:accuracy + vividness + block:vividness + accuracy:vividness + block:accuracy:vividness + (1 + block \| subject) + (1 + block + accuracy + block:accuracy + vividness + block:vividness + accuracy:vividness + block:accuracy:vividness \| connection) |

**Table S5**. Model structures for connectivity analysis of imCOH, Granger Causality, and time-reversed Granger Causality. The weight or probability of the model is computed with reference to the summed probability of the models that share the same null model and the corresponding null model itself. Full statistical report is provided as a separate dataset.

| Local & long-range PAC (binomial & Gaussian) | | |
|---|---|---|
| **model** | **Null** | **Formula** |
| intercept | - | PAC ~ 1 + (1 \| subject) + (1 \| region or pair of regions) |
| period | intercept | PAC ~ 1 + period + (1 \| subject) + (1 \| region or pair of regions) |
| period rnd1 | intercept | PAC ~ 1 + period + (1 \| subject) + (1 + period \| region or pair of regions) |
| period rnd2 | intercept | PAC ~ 1 + period + (1 + period\| subject) + (1 + period \| region or pair of regions) |
| block | period rnd2 | PAC ~ 1 + period + block + (1 + period\| subject) + (1 + period \| region or pair of regions) |
| block rnd1 | period rnd2 | PAC ~ 1 + period + block + (1 \| subject) + (1 + block \| region or pair of regions) |
| block rnd2 | period rnd2 | PAC ~ 1 + period + block + (1 + block\| subject) + (1 + block \| region or pair of regions) |
| full | block rnd2 | PAC ~ 1 + period + block + period:block + (1 + block\| subject) + (1 + block \| region or pair of regions) |
| full rnd 1 | block rnd2 | PAC ~ 1 + period + block + period:block + (1 + block\| subject) + (1 + block + period:block \| region or pair of regions) |
| full rnd 2 | block rnd2 | PAC ~ 1 + period + block + period:block + (1 + block + period:block \| subject) + (1 + block + period:block \| region or pair of regions) |

**Table S6.** Model structures for phase-amplitude coupling (PAC) statistical analysis. The same models were used for local and long-range PAC and for proportions of significant channels or pairs (binomial) and PAC strength (Gaussian). The weight or probability of the model is computed with reference to the summed probability of the models that share the same null model and the corresponding null model itself. Full statistical report is provided as a separate dataset.

| Local & long-range PAC (binomial & Gaussian) | | |
|---|---|---|
| **model** | **null** | **Formula** |
| intercept | - | PAC ~ 1 + (1 \| subject) + (1 \| region) |
| block | intercept | PAC ~ 1 + block + (1 \| subject) + (1 \| region) |
| block rnd1 | intercept | PAC ~ 1 + block + (1 \| subject) + (1 + block \| region) |
| block rnd2 | intercept | PAC ~ 1 + block + (1 + block\| subject) + (1 + block \| region) |
| vividness | block rnd2 | PAC ~ 1 + block + vividness + (1 + block\| subject) + (1 + block \| region) |
| vividness rnd1 | block rnd2 | PAC ~ 1 + block + vividness + (1 + block\| subject) + (1 + block + vividness \| region) |
| vividness rnd2 | block rnd2 | PAC ~ 1 + block + vividness + (1 + block + vividness\| subject) + (1 + block + vividness \| region) |
| full | vividness rnd2 | PAC ~ 1 + block + vividness + block:vividness + (1 + block + vividness\| subject) + (1 + block + vividness \| region) |
| full rnd 1 | vividness rnd2 | PAC ~ 1 + block + vividness + block:vividness + (1 + block + vividness \| subject) + (1 + block + vividness + block:vividness \| region) |
| full rnd 2 | vividness rnd2 | PAC ~ 1 + block + vividness + block:vividness + (1 + block + vividness + block:vividness \| subject) + (1 + block + vividness + block:vividness \| region) |

**Table S7.** Model structures for secondary phase-amplitude coupling (PAC) statistical analysis including vividness as a predictor. The models were applied to the delay period only. The same models were used for local and long-range PAC and for proportions of significant channels or pairs (binomial) and PAC strength (Gaussian). The weight or probability of the model is computed with reference to the summed probability of the models that share the same null model and the corresponding null model itself. Full statistical report is provided as a separate dataset.

| Local & long-range PAC (binomial & Gaussian) | | |
|---|---|---|
| **model** | **null** | **Formula** |
| intercept | - | PAC ~ 1 + (1 \| subject) + (1 \| region) |
| block | intercept | PAC ~ 1 + block + (1 \| subject) + (1 \| region) |
| block rnd1 | intercept | PAC ~ 1 + block + (1 \| subject) + (1 + block \| region) |
| block rnd2 | intercept | PAC ~ 1 + block + (1 + block\| subject) + (1 + block \| region) |
| accuracy | block rnd2 | PAC ~ 1 + block + accuracy + (1 + block\| subject) + (1 + block \| region) |
| accuracy rnd1 | block rnd2 | PAC ~ 1 + block + accuracy + (1 + block\| subject) + (1 + block + accuracy \| region) |
| accuracy rnd2 | block rnd2 | PAC ~ 1 + block + accuracy + (1 + block + accuracy \| subject) + (1 + block + accuracy \| region) |
| full | accuracy rnd2 | PAC ~ 1 + block + accuracy + block:vividness + (1 + block + accuracy \| subject) + (1 + block + accuracy \| region) |
| full rnd 1 | accuracy rnd2 | PAC ~ 1 + block + accuracy + block:vividness + (1 + block + accuracy \| subject) + (1 + block + accuracy + block: accuracy \| region) |
| full rnd 2 | accuracy rnd2 | PAC ~ 1 + block + accuracy + block:vividness + (1 + block + accuracy + block: accuracy \| subject) + (1 + block + accuracy + block: accuracy \| region) |

**Table S8.** Model structures for secondary phase-amplitude coupling (PAC) statistical analysis including accuracy as a predictor. The models were applied to the delay period only. The same models were used for local and long-range PAC and for proportions of significant channels or pairs (binomial) and PAC strength (Gaussian). The weight or probability of the model is computed with reference to the summed probability of the models that share the same null model and the corresponding null model itself. Full statistical report is provided as a separate dataset.
